# Systematic characterization of *Ustilago maydis* sirtuins shows Sir2 as a modulator of pathogenic gene expression

**DOI:** 10.1101/2023.02.28.530406

**Authors:** Blanca Navarrete, José I. Ibeas, Ramón R. Barrales

## Abstract

Phytopathogenic fungi must adapt to the different environmental conditions found during infection and avoid the immune response of the plant. For these adaptations, fungi must tightly control gene expression, allowing sequential changes in transcriptional programs. In addition to transcription factors, chromatin modification is used by eukaryotic cells as a different layer of transcriptional control. Specifically, the acetylation of histones is one of the chromatin modifications with a strong impact on gene expression. Hyperacetylated regions usually correlate with high transcription and hypoacetylated areas with low transcription. Thus, histone deacetylases (HDACs) commonly act as repressors of transcription. One member of the family of HDACs is represented by sirtuins, which are deacetylases dependent on NAD+, and, thus, their activity is considered to be related to the physiological stage of the cells. This property makes sirtuins good regulators during environmental changes. However, only a few examples exist, and with differences in the extent of the implication of the role of sirtuins during fungal phytopathogenesis. In this work, we have performed a systematic study of sirtuins in the maize pathogen *Ustilago maydis*, finding Sir2 to be involved in the dimorphic switch from yeast cell to filament and pathogenic development. Specifically, the deletion of *sir2* promotes filamentation, whereas its overexpression highly reduces tumor formation in the plant. Moreover, transcriptomic analysis revealed that Sir2 represses genes that are expressed during biotrophism development. Interestingly, our results suggest that this repressive effect is not through histone deacetylation, indicating a different target of Sir2 in this fungus.

## Introduction

Phytopathogenic fungi must sense many environmental host cues and respond with developmental changes in order to ensure proper plant infection progression. In the well-established model organism *Ustilago maydis*, a biotrophic pathogen infecting maize plants, the first step in the pathogenic program is the switch from yeast to filament on the surface of the plant, followed by the fusion of two sexually compatible filaments. This dikaryotic filament blocks the cell cycle and extends until it identifies the appropriate location for penetration, where it develops a specialized invasive structure, the appressoria. Upon plant penetration, the filament releases the cell cycle block and colonizes the plant until the development of teliospores inside plant tumors induced by the fungus (Brefort et al., 2009; Vollmeister et al., 2012; Castanheira and Pérez-Martín, 2015). All these sequential changes must be tightly controlled in order to ensure successful infection. Many advances have been achieved in determining the transcription factors involved in this control mechanism in many fungi. In *U. maydis*, environmental cues are transmitted by the MAP kinases and cAMP signaling pathways to the Prf1 transcription factor, which controls, among others, the bE/bW compatible heterodimer transcribed from the MAT b locus in the dikaryon. This heterodimer controls many downstream virulence genes, including other important transcription factors (Hartmann et al., 1996, Hartman et al., 1999; Brachmann et al., 2001; Heimel et al., 2010; Lanver et al., 2017).

Another layer of control is chromatin modification, which plays crucial roles in transcriptional regulation in response to environmental cues (Mazzio and Soliman, 2012; Rando and Winston, 2012; Badeaux and Shi, 2013). Histone modifiers carry out different posttranslational modifications in the histone tails, such as acetylation, methylation or phosphorylation, among others, with potential alterations in the transcriptional stages of the surrounding area. One of the major and most well-described histone modifications, together with methylation, is the acetylation in a lysine residue (Allfrey et al., 1964). The enzymes responsible for this acetylation are called histone acetyltransferases (HATs), and those involved in the removal of this modification are the histone deacetylases (HDACs).

The role of these chromatin modifiers in pathogenesis has been explored in different fungi. Gcn5 is the main HAT studied in plant pathogenic fungi, with important roles in development and infection (Ruijter et al., 2003; González-Prieto et al., 2014; Kong et al., 2018; Liu et al., 2022). In *U. maydis*, the deletion of Gcn5 causes constitutive filamentation and reduction of infection by, at least, the derepression of *prf1* and *bE1* genes (González-Prieto et al., 2014). On the other hand, the roles of HDACs during fungal plant pathogenesis, although poorly characterized, are better known than those for HATs. HDACs can be classified into three main different categories, Class I, II and III, based on their homology to the yeast orthologues Rpd3, Hda1 and Sir2, respectively (Ruijter et al., 2003). Among Class I/II HDACs, the Set3 complex, comprising the HDAC Hos2, is one of the best studied in fungal plant pathogens, with conserved roles in pathogenesis (Elías-Villalobos et al., 2019). In *U. maydis*, Hos2 affects filamentation and pathogenesis through direct regulation of the MAT a locus (Elías-Villalobos et al., 2015). In addition to Hos2, other Class I/II HDACs, Hda1 and Hda2, have also been characterized in *U. maydis*. Hda1 is essential for teliospore production with a role in gene regulation, repressing the transcription of *egl1* and *mig1* during the non-pathogenic state of the fungus (Reichmann et al., 2002; Torreblanca et al., 2003). In contrast, deletion of *hda2* did not alter the infection capability of *U. maydis*. (González-Prieto et al., 2004; Elías-Villalobos et al., 2015). Class III HDACs constitute a particular group of histone deacetylases that are dependent on NAD^+^ for their catalytic activity (Tanner et al., 2000; Tanny and Moazed, 2001; Jackson and Denu, 2002; Zhao and Rusche, 2022). The founding member of this class, collectively named sirtuins, is Sir2 (Silent Information Regulator 2) from *Saccharomyces cerevisiae*. ScSir2 forms a complex with other SIR proteins and is involved in the silencing of heterochromatin-like regions in this yeast by deacetylating H4 lysine 16 residue (H4K16) (Robyr et al., 2002; Suka et al., 2002). The role of Sir2 in chromatin silencing is broadly observed, with examples also described in filamentous fungi, such as *Neurospora crassa* or *Aspergillus nidulans* (Smith et al., 2008; Shimizu et al., 2012; Itoh et al., 2017), which suggests an ancient role of this protein in silencing (Hickman et al., 2011). In addition to Sir2, other sirtuins have been characterized in different organisms. In *S. cerevisiae* and *Schizosaccharomyces pombe*, all the other sirtuins, Hst1 to 4 in *S. cerevisiae* and Hst2 and 4 in *S. pombe*, have been linked to chromatin silencing as well as direct gene regulation (Brachmann et al., 1995; Freeman-Cook et al., 1999; Sutton et al., 2001; Halme et al., 2004; Wilkins et al., 2014). Within the sirtuin family, Sir2 has been described to control pathogenesis in different fungi. For instance, in the human pathogen *Candida glabrata*, Sir2 represses the EPA adhesin genes, which are essential for infection (Domergue et al., 2005), and in *Cryptococcus neoformans*, Sir2 is essential for virulence, due to a mechanism not described so far (Arras et al., 2017). The main example to date for the role of Sir2 in plant pathogens is found in *Magnaporthe oryzae*. In this rice pathogen, Sir2 likely affects infection through inactivation by deacetylation of the MoJmjC repressor, which would lead to an increase in superoxide dismutase expression, allowing ROS detoxification (Fernandez et al., 2014).

In order to increase our knowledge regarding the role of sirtuins in fungal plant pathogens, we have performed a characterization of the sirtuin family in *U. maydis*. We have observed that two of the five sirtuins present in *U. maydis*, Sir2 and Hst4, display nuclear localization during the entire cell cycle. From them, Hst4 is essential and Sir2 negatively impacts the yeast to filament transition and virulence. While the deletion of *sir2* slightly increases the virulence capacity, its overexpression significantly reduces virulence. A transcriptomic analysis of both deletion in filamentation conditions and overexpression during infection indicates that Sir2 avoids the proper activation of a group of genes induced during the biotrophic development. We have observed an increase in H4 acetylation in a Δ*sir2* mutant in the upregulated genes. However, this deacetylation is not detected in the typical residue observed in other organisms, lysine 16. As these effects may be the consequence of the increase in transcription observed, further analyses are required in order to detect the specific target of Sir2 in terms of its regulatory role in *U. maydis*.

## Materials and Methods

### Strains and growth condition

*Escherichia coli* DH5α, pJET1.2/blunt (Thermo Scientific, Carlsbad, CA, USA) and pBluescript II SK (+) (Stratagene, San Diego, CA, USA) were used for cloning purposes. The growth conditions for *E. coli* were described in (Sambrook et al., 1989). All the strains used in this study are derived from the haploid pathogenic SG200 strain and are listed in Supplementary Table S1. As previously described in (Gillissen et al., 1992), *U. maydis* cultures were performed in YEPSL (0.4% bactopeptone, 1% yeast extract and 0.4% saccharose) at 28ºC, unless otherwise specified. For charcoal filamentation assays, exponential cultures were spotted onto PD–charcoal plates (2.4% potato dextrose broth, 1% charcoal, 2% agar) and grown for 18-20 hours at 25ºC. Pathogenicity assays were performed as described in (Kämper et al., 2006). *U. maydis* exponential cultures were concentrated to an OD_600_ of 0.5 or 1 and injected into 7-day-old maize (*Zea mays*) seedlings (Early Golden Bantam). Disease symptoms were quantified at 14 dpi. Data of individual infection experiments are listed in Supplementary Table S2. Statistical analyses were performed in the GraphPad Prism 8 software.

### Plasmid and strain construction

Molecular biology techniques were used, as previously described (Sambrook et al., 1989). *U. maydis* DNA isolation and transformation procedures were carried out following the protocol described in (Schulz et al., 1990). Deletion mutants for *sir2* (UMAG_00963), *hst2* (UMAG_05892), *hst4* (UMAG_05758) and *hst5* (UMAG_05239) were generated by homologous recombination, as described previously (Brachmann et al., 2004; Kämper, 2004). Deletion for *hst6* (UMAG_12006) was performed using the NEBuilder HiFi DNA Assembly (New England Biolabs, Ipswich, MA, USA) system. The *sir2* complementation mutant was generated by reintroducing the *sir2* open reading frame (ORF) in the Δ*sir2* background in its endogenous loci, replacing the nat resistance cassette of the Δ*sir2* mutant with the *sir2* ORF, followed by the geneticin resistance cassette by homologous recombination. For *sir2* overexpression with the *otef* promoter, the p123-P*otef*:*sir2* plasmid was generated by replacing the eGFP fragment from the p123 plasmid (Aichinger et al., 2003) with the *sir2* ORF. The *sir2* ORF was amplified by PCR using Q5 High-Fidelity DNA polymerase (New England Biolabs, Ipswich, MA, USA) and cloned into p123 within the NcoI and NotI restriction sites of p123. For *sir2* overexpression with the *pit2* promoter, we constructed the p123-P*pit2*:*sir2* plasmid. The *sir2* ORF was amplified and cloned into p123-P*pit2* within the NcoI and XbaI restriction sites of p123-P*pit2*. To generate SG200 P*otef*:*sir2* or SG200 P*pit2*:*sir2*, p123-P*otef*:*sir2* or p123-P*pit2*:*sir2* was linearized with SspI and integrated into the *ip* locus by homologous recombination. For GFP endogenous sirtuin tagging, the plasmids pBSK-*sir2*:eGFP, pBSK-*hst2*:eGFP, pBSK-*hst4*:eGFP, pBSK-*hst5*:eGFP and pBSK-*hst6*:eGFP were generated using the NEBuilder HiFi DNA Assembly (New England Biolabs, Ipswich, MA, USA) system. A 1 Kb fragment containing the gene of interest (ORF without the STOP codon) and a 1 Kb fragment of the 3’ region were amplified using primers designed in the NEBuilder assembly tool. eGFP followed by hygromycin resistance cassettes were amplified from the pmf5-1h plasmid (Becht et al., 2006). All the fragments were cloned into the pBluescript II SK (+) plasmid using the NEBuilder HiFi DNA Assembly (New England Biolabs, Ipswich, MA, USA). Constructs were amplified by PCR prior to their transformation in *U. maydis*. The primers used in this study are listed in Supplementary Table S3. All the strains and numbers of copies integrated into the *ip* locus were verified by PCR and Southern blotting.

### Microscopy and image analysis

To analyze the filamentation capability of *U. maydis* in PD-charcoal plates, single colonies where visualized using the Leica M205 Stereoscope equipped with an ORCA-Flash4.0 LT Hamamatsu digital camera. The area of the colonies was measured by selecting the perimeter of each colony using the plugging convex hull of ImageJ software. To determine sirtuins’ localization, *sir2*:eGFP, *hst2*:eGFP, *hst4*:eGFP, *hst5*:eGFP and *hst6*:eGFP cells were visualized using a DeltaVision microscopy system comprising an Olympus IX71 microscope and CoolSnap HQ camera (Applied Precision, Issaquah WA, USA). To visualize mitochondria, 0.5 mM Mito-Tracker CM-H2Xros (Molecular Probes, Eugene, OR) was added to the *U. maydis* YEPSL cell culture and cells were incubated for 15 min at 25ºC (Bortfeld et al., 2004). To analyze the *U. maydis* progression inside the maize plant, leaves samples from 3, 4 and 6 dpi infected plants were distained with ethanol, treated for 4 h at 60ºC with 10% KOH, washed in phosphate buffer and then stained with propidium iodide (PI) to visualize plant tissues in red and wheat germ agglutinin (WGA)/AF488 to visualize the fungus in green. At least four leaves from two independent experiments were stained and visualized by fluorescence microscopy (Leica SPE DM2500, Leica, WZ, Germany). Image processing was carried out using the ImageJ software.

### RNA extraction and RT-qPCR

Total RNA was extracted from *U. maydis* cells grown in YEPSL medium, PD–charcoal plates and from infected leaves by excising 2–3 cm segments from below the injection holes. All the samples were ground into a powder using a mortar/pestle under liquid nitrogen. Total RNA was purified using TRIzol reagent (Invitrogen, Carlsbad, CA, USA) and the Direct-zol RNA Miniprep Plus Kit (Zymo Research, Irvine, CA, USA). RNA was retrotranscribed from 3 µg of total RNA using the RevertAid H Minus First Strand cDNA Synthesis Kit (Thermo Scientific, Carlsbad, CA, USA). RT-qPCR was performed using a Real-Time CFX Connect (Bio-Rad, Hercules, CA, USA) and SYBR Premix Ex Taq II (Tli RNase H Plus) (Takara Bio INC, Kusatsu, Japan). All reactions were performed in at least three biological replicates, and gene expression levels were calculated relative to the expression levels of the constitutively expressed fungal *ppi1* gene. Primers used for RT-qPCR are listed in Supplementary Table S2. The quantification of relative fungal biomass in infected maize leaves was performed as previously described (Brefort et al., 2014). *U. maydis* biomass was quantified measuring the signal of the *ppi1* fungal gene relative to the plant gene GAPDH. Statistical analyses were performed in the GraphPad Prism 8 software.

### RNA-Seq analysis

Total RNA extracted from axenic cultures and PD–charcoal plates from *U. maydis* wild-type and Δ*sir2* strains was submitted to BGI TECH SOLUTIONS (HONGKONG) CO., LIMITED, in a 200-500 ng/μl concentration, with a total RNA quantity of 5 - 8 μg and quality parameters of OD_260/280_ = 1.8-2.1 and OD_260/230_ > 1.5. The BGI company prepared all libraries and performed the single-end sequencing via the BGISEQ-500 RNA-Seq service. Two replicates of each strain and condition were processed. To determine Sir2-regulated genes during pathogenesis, 7-day-old maize seedlings were infected with wild-type and P*pit2*:*sir2* >1c strains and total RNA was purified at 3 dpi. RNA samples were submitted to BGI TECH SOLUTIONS (HONGKONG) CO., LIMITED, and paired-end sequenced via the DNBseq PE100 service. Three replicates of each strain were processed. Reads were mapped to the *U. maydis* genome using HISAT2 and reads from infected plant tissues were previously filtered against the annotated maize genome. Reads were counted for *U. maydis* using the HTseq tool in the Galaxy platform, and, for expression analysis, only uniquely mapping exon read counts were considered. Pairwise comparisons were performed using the R package DESeq2 (Love et al., 2014). Genes with log2 fold change > 0.5 or <-0.5 and adjusted p-value < 0.05 were considered differentially regulated.

### Western blot analysis

For total protein extraction, cells from exponential culture were collected by centrifugation and washed twice with 20 mM Tris–HCl pH 8.8. Pellets were ground to powder with a mortar under liquid nitrogen and resuspended in RIPA buffer (50 mM Tris–HCl, pH 8, 150 mM NaCl, 1% Nonidet P-40, 0.5% sodium deoxycholate, 0.1% SDS) supplemented with 1 μg/ml Pepstatin A (PanReac AppliChem, Barcelona, Spain), 1 μg/ml Bestatin (Thermo Scientific, Carlsbad, CA, USA), 1mM PMSF (PanReac AppliChem, Barcelona, Spain) and EDTA-free protease inhibitor complex (cOmplete Tablets EDTA-free, Roche, Mannheim, BW, Germany). After cell lysis, samples were centrifuged and the supernatant was collected. For protein extraction in PD–charcoal plates, cells were scraped off and ground to powder in liquid nitrogen and resuspended in 12% TCA solution to precipitate proteins. Pellets were washed 4 times with ice-cold acetone and dissolved in extraction buffer (100 mM Tris–HCl pH 8, 50 mM NaCl, 1% SDS, 1mM EDTA) supplemented with the protease inhibitors listed above. Protein concentration was measured by the BCA protein assay. Here, 40 μg of protein extract was loaded into a 10% TGX Stain-Free FastCast Acrylamide Gel (Bio-Rad, Hercules, CA, USA) or SDS polyacrylamide 15% running gel in the case of histone analysis. Separated proteins were transferred into a nitrocellulose membrane using the Trans-Blot Turbo transfer system (Bio-Rad, Hercules, CA, USA). The membrane was incubated with mouse polyclonal anti-GFP antibody (Roche, Mannheim, BW, Germany) (1:1000 in PBST 5% fat-free milk) and anti-mouse IgG HRP conjugated (Invitrogen, Carlsbad, CA, USA) (1:5000) was used as a secondary antibody. Histone modifications were detected with primary antibodies specific to H3 (Sigma-Aldrich, Darmstadt, Germany), H3ac (Sigma-Aldrich, DA, Germany), H3K9ac (abcam, Cambridge, UK) (1:5000 in PBST 5% fat-free milk), H4 (Sigma-Aldrich, Darmstadt, Germany), H4ac (Sigma-Aldrich, Darmstadt, Germany) (1:5000 in PBST 3% BSA) and H4K16ac (Sigma-Aldrich, Darmstadt, Germany) (1:5000 in PBST 5% fat-free milk) and anti-rabbit HRP conjugated as a secondary antibody (Sigma-Aldrich, Darmstadt, Germany) (1:5000). Chemiluminescence was detected with SuperSignal™ West Femto Maximum Sensitivity Substrate (Thermo Scientific, Carlsbad, CA, USA). Image gel and membrane acquisition was carried out with the ChemiDoc XRS (Bio-Rad, Hercules, CA, USA). All the Western blot assays were performed with at least three biological replicates and quantified using the Image Lab software.

### Chromatin Immunoprecipitation (ChIP)

Exponential culture of *U. maydis* cells were cross-linked by incubating with 1% formaldehyde for 10 minutes and reaction stopped by adding glycine to a final concentration of 250 mM for 10 minutes at room temperature. Cells were collected by centrifugation and washed twice with cold PBS. 2x 250 mg of pellets were ground to powder with a mortar under liquid nitrogen and resuspended in ChIP lysis Buffer (50 mM HEPES-KOH pH 7.5, 140 mM NaCl, 1 mM EDTA pH 8, 1% Triton X-100, 0.1% Na-deoxycholate, 0.1% SDS) supplemented with 1 μg/ml Pepstatin A (PanReac AppliChem, Barcelona, Spain), 1 μg/ml Bestatin (Thermo Scientific, Carlsbad, CA, USA), 1mM PMSF (PanReac AppliChem, Barcelona, Spain) and EDTA-free protease inhibitor complex (cOmplete Tablets EDTA-free, Roche, Mannheim, BW, Germany). Samples were then sonicated in a Bioruptor® sonication device (Diagenode) for 20 min, with 2 minutes pulses separated by 1 minute rest periods at maximum power. 100 μl of the chromatin extract was kept as input and a total of 10 O.D. of chromatin extract was used for IP. Samples were incubated with 3 μl of antibodies against H3ac (Sigma-Aldrich, DA, Germany), H4 (Sigma-Aldrich, Darmstadt, Germany), H4ac (Sigma-Aldrich, Darmstadt, Germany) and H4K16ac (Sigma-Aldrich, Darmstadt, Germany) at 4°C overnight on a rotary shaker. Precipitation of the protein-antibody conjugate was performed incubating with Dynabeads® Protein A (Thermo Scientific, Carlsbad, CA, USA) 40 minutes at 4ºC in a rotary shaker. Beads were washed twice with WB150 (20 mM Tris-HCl pH 8, 150 mM NaCl, 2 mM EDTA pH 8, 1% Triton X-100), once with WB500 (20 mM Tris-HCl pH 8, 500 mM NaCl, 2 mM EDTA pH 8, 1% Triton X-100) and eluted in TES buffer (50 mM Tris-HCl pH 8, 10 mM EDTA, 1% SDS). To reverse the crosslink, both input and IP chromatin extracts were incubated at 65°C for 16 hours. Histone-bound DNA was treated with Proteinase K (Thermo Scientific, Carlsbad, CA, USA) and DNA purification was done using the ChIP DNA Clean & Concentrator™ (Zymo Research, Irvine, CA, USA). For RT-qPCR a 20-fold dilution of each immunoprecipitated sample and a 200-fold dilution of input samples were used. Primers used for each amplicon were listed in Supplementary Table S2. All experiments were performed with three biological replicates.

## Results

### The systematic characterization of sirtuins in *U. maydis* shows Sir2 as a nuclear sirtuin controlling cell-to-filament transition

In a previous phylogenetic analysis (Elías-Villalobos et al., 2019), five sirtuin homologs were described in *U. maydis*: UMAG_00963 (Sir2), UMAG_05892 (Hst2), UMAG_05758 (Hst4), UMAG_05239 (Hst5) and UMAG_12006 (Hst6). As expected, all these proteins contained the conserved sirtuin catalytic domain (PROSITE:PS50305) involved in protein deacetylation (Figure 1A). Additionally, we detected nuclear localization signals in only Sir2 and Hst4 (Figure 1A). As we were interested in studying the possible role of sirtuins in the control of the transcriptional pathogenic program, we examined the cellular localization of these proteins in order to focus on the nuclear ones. Consistent with their localization motifs, we observed that Sir2 and Hst4 displayed nuclear localization (Figure 1B). Additionally, Hst2 showed a nuclear signal only in mitotic cells (Figure 1B). By contrast, Hst5 and 6 were localized in the mitochondria, as verified by Mito-tracker colocalization (Figure 1B and Supplementary Figure S1A). This is consistent with the similarity described for these sirtuins to mitochondrial ones (Elías-Villalobos et al., 2019). In addition to the cellular localization study, we deleted all sirtuin genes in the solopathogenic strain SG200, except for the *hst4* gene, whose deletion we found to be lethal (Supplementary Figure S2B). We performed plant infection assays with these mutants and found no significant changes in the symptoms of plants infected with *∆hst2, ∆hst5* or *∆hst6* mutants compared to the wild-type strain (Supplementary Figure S1C). However, we detected a slight increase in the symptoms of plants infected with the *∆sir2* mutant (Figure 1C). A more significant difference was observed when we studied the yeast to filament transition in solid PD–charcoal plates, which mimicked the hydrophobic conditions of the plant surface. Here, we observed an increase in the filamentation capability of the *∆sir2* mutant (Figure 1D and Supplementary Figure S1D), which was restored to its normal level after the reintroduction of *sir2* in the endogenous locus (Supplementary Figure S1D). We quantified this increase in filamentation by growing individual colonies on PD-charcoal plates and measuring filaments length of the single colonies, observing that *∆sir2* colonies exhibit longer filaments compared to wild-type (Figure 1D). No changes in filamentation were observed for the rest of the sirtuin mutants (Supplementary Figure S1D).

**FIGURE 1.**
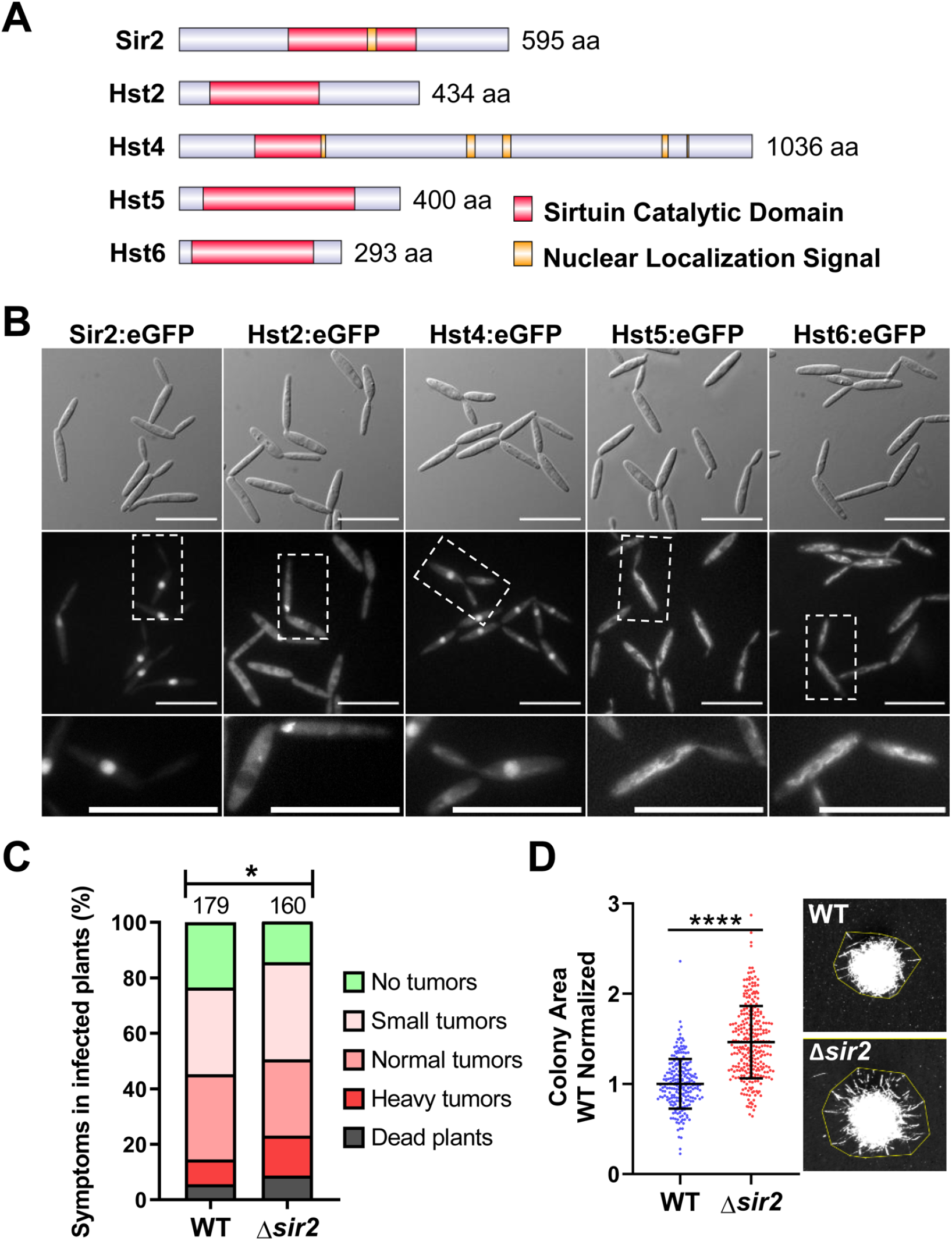
The nuclear sirtuin Sir2 is involved in virulence and filamentation of *Ustilago maydis*. **(A)** Schematic representation of all the sirtuins identified in *U. maydis*. **(B)** Subcellular localization of the indicated *U. maydis* sirtuins tagged with eGFP in its endogenous *loci*. Scale bar represents 20 μm. **(C)** Quantification of symptoms for plants infected with the wild-type and Δ*sir2* mutant at 14 dpi. Total number of infected plants is indicated above each column. Three biological replicates were analyzed. Mann–Whitney statistical test was performed (* p-value < 0.05). **(D)** Quantification of the area of the wild-type and Δ*sir2* mutant single colonies grown on PD–charcoal plates for 48 hours at 25ºC. The colony area was measured as indicated in the stereoscopic images. Data was normalized with the mean of the area of the wild-type colonies. Three biological replicates were analyzed. Student’s t-test statistical analysis was performed (**** p-value < 0.001).

### Sir2 affects the transcription of some genes during filamentation

In order to disregard a possible pleiotropic effect of *sir2* deletion, we performed growth assays in rich (YEPSL), complete (CMD) and minimal (MMD) media (Supplementary Figure S2A), flow cytometry analysis of DNA-stained cells (Supplementary Figure S2B) and a stress assay with sorbitol and NaCl as osmotic stressors, H_2_O_2_ as an oxidant, SDS as a membrane-perturbing drug, calcofluor white (CFW) and Congo red as cell wall integrity sensors and Tunicamycin and DTT as endoplasmic reticulum stressors (Supplementary Figure S2C). We were not able to detect any significant differences with respect to wild-type cells, suggesting no pleiotropy. Thus, we focused on the role of Sir2 in cell to filament transition. We analyzed Sir2 protein levels and observed a drastic reduction in Sir2 during filamentation (Figure 2A). Although Sir2 is mainly present in axenic conditions, and its deletion caused an increase in filamentation on PD–charcoal plates, no filaments were found in the *∆sir2* mutant in axenic conditions (Figure 2B). In addition, almost no differences in gene expression were found by RNA sequencing (RNA-seq) analysis in a *∆sir2* mutant in axenic conditions (Figure 2C and Supplementary Table S4). The most upregulated gene was *eff1-9* (Figure 2C and Supplementary Table S4), a member of the *eff* family of effector proteins important for virulence (Khrunyk et al., 2010; Schuster et al., 2018). *eff1-9* upregulation was confirmed by RT-qPCR in the *∆sir2* mutant and the transcription levels were restored in the complementation strain (Supplementary Figure S2D). The other upregulated genes were two subtelomeric genes, UMAG_04104 and UMAG_05781, and three genes involved in metabolism, UMAG_01476, UMAG_04922 and UMAG_01656 (Figure 2C and Supplementary Table S4). We then analyzed transcription changes in a *∆sir2* mutant in filament-induced PD–charcoal plates by RNA-seq. We obtained 11 downregulated and 31 upregulated genes (Figure 2D and Supplementary Table S5). Many of the upregulated genes, 58%, encoded for predicted effector proteins, including many of the previously characterized ones: Mig2-6 (Farfsing et al., 2005), Pit1 and 2 (Doehlemann et al., 2011; Mueller et al., 2013), Eff1-7 (Khrunyk et al., 2010), Cmu1 (Djamei et al., 2011), Rsp3 (Ma et al., 2018), Erc1 (Ökmen et al., 2022) and Egl1 and 3 (Schauwecker et al., 1995; Doehlemann et al., 2008). In order to know if the *∆sir2* upregulated genes are genes which have to be expressed when filamentation is induced in PD-charcoal plates, we studied the distribution of the significantly different log2 fold changes in expression of all these genes during charcoal growth, in comparison with axenic conditions, in a wild-type strain (Supplementary Table S6). We observed that when all genes of a wild-type strain were analyzed, there was a general distribution in which log2 fold changes expanded from negative to positive values (Figure 2E). However, the group of genes that were upregulated in *∆sir2* corresponded to genes upregulated during charcoal growth (Figure 2E, Supplementary Table S5 and Supplementary Table S6). As comparison between growth on charcoal plates versus liquid rich media imply other changes different than filamentation, we crossed our data with other datasets more specific for filamentation. Interestingly, we found that 42% of the upregulated genes in the Δ*sir2* mutant are genes upregulated when filamentation and appressoria formation is induced by an hydrophobic surface and hydroxy fatty acids (Lanver et al., 2014) (Supplementary Table S5). Interestingly, *egl1* has been previously identified as a gene specifically expressed in filaments (Schauwecker et al., 1995). As many of the described effectors found to be upregulated in a Δ*sir2* mutant have their effect during infection, we also studied the possibility of these genes being activated during the infection process. In a previous study of the transcriptional changes observed during infection performed by RNA-seq, Lanver *et al*., 2018 (Lanver et al., 2018) described different modules of coexpressed genes during infection. We observed the strong enrichment of genes belonging to the cyan (26%) and the magenta (52%) modules in the group of genes upregulated in *∆sir2* in comparison to all the upregulated genes during filamentation (Figure 2F Supplementary Table S5 and Supplementary Table S6). The magenta module correlates with the establishment and maintenance of biotrophy, while the cyan one represents a tumor module (Lanver et al., 2018). All these data may indicate a role of Sir2 in avoiding the proper activation of a group of genes induced during filamentation and probably during infection.

**FIGURE 2.**
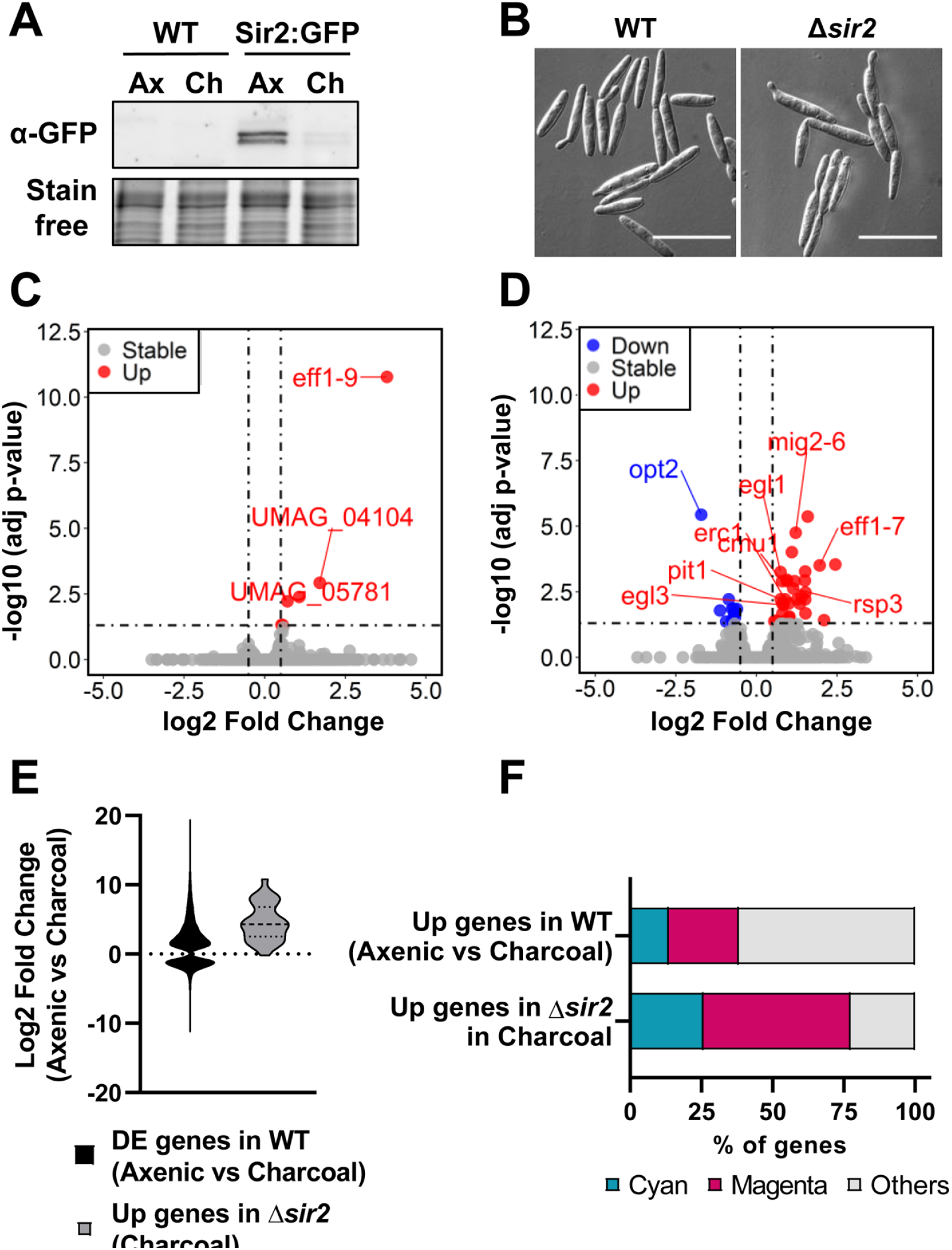
Sir2 is degraded during filamentation and is involved in the repression of a group of filamentation-induced genes. **(A)** Western blot showing Sir2:eGFP protein levels of wild-type strain. Total proteins were extracted from cells growing in axenic culture (Ax) and PD–charcoal plates (Ch) for 18 hours at 25ºC. Stain-free gel is shown as a loading control. **(B)** Images of wild-type and Δ*sir2* mutant growing in axenic culture. Scale bar represents 20 μm. **(C, D)** Volcano plot showing the log2 fold change in gene expression and the statistical significance of the differential expression analysis from RNA-seq data obtained for the Δ*sir2* mutant compared to wild-type in axenic culture **(C)** and PD–charcoal plates **(D)**. Red, blue and grey dots represent the upregulated genes (log2 fold change ≥ 0.5, adjusted p-value < 0.05), downregulated genes (log2 fold change ≤ −0.5 adjusted p-value < 0.05) and genes without changes in the Δ*sir2* mutant, respectively. *sir2* data has been removed for plotting purpose. **(E)** Log2 fold change distribution of the differentially expressed genes (adjusted p-value < 0.05) for wild-type in PD–charcoal plates compared to axenic culture (green) and for the upregulated genes in the Δ*sir2* mutant compared to wild-type in PD–charcoal plates (red). **(F)** Percentage of genes belonging to the magenta or cyan modules of coexpressed genes during infection for the upregulated genes of wild-type in PD–charcoal plates compared to axenic culture and for the upregulated genes in the Δ*sir2* mutant compared to wild-type in PD–charcoal plates.

### Sir2 overexpression drastically reduces infection capability

The observation that Sir2 affects the expression of genes that are induced during infection led us to study the role of Sir2 during plant infection. As the deletion of *sir2* showed a slight increase in infected plant symptoms (Figure 1C), and Sir2 seemed to repress genes activated during infection (Figure 2F), we decided to study the effect of *sir2* overexpression. First, we integrated one or more than one copy of the *sir2* gene under the control of the constitutive *otef* promoter in the *ip* locus. No growth defects were detected in the overexpression mutant (Supplementary Figure S3A), however, as it can be observed in Figures 3A, B and Supplementary Figure S3C, the filamentation capability was reduced according to the *sir2* expression level found in both mutants. These data confirm the role of Sir2 in avoiding the proper induction of the filamentation process. Additionally, we overexpressed *sir2* during pathogenesis by using the *pit2* promoter, which reaches its expression peak at 2 days post-infection (dpi) (Figure 4C). We infected maize plants with mutants harboring one or more than one copy of *Ppit2:sir2*, which did not show any significant defect in growth in different tested media (Supplementary Figure S3B), and quantified the symptoms in infected plants. The size of tumors was clearly reduced when *sir2* was significantly induced at 3 dpi (Figures 3C, D), indicating that Sir2 affects the infection process during plant colonization. When we analyzed the pathogenic defects of this mutant, we did not detect any alteration in fungi morphology during plant colonization (Figure 4A). However, after 3 dpi, we detected a gradual reduction in fungal biomass during the progression of infection (Figure 4B). It is interesting to note that the reduction in fungal biomass was detected several days after the main overexpression of *sir2*, obtained at 2 dpi (Figure 4C). A possible explanation is that the overexpression of *sir2* during the biotrophy establishment (2 dpi), affects the transcription of effector genes essential for this process or genes with roles from 4 dpi onward. Another alternative may be that the reduction in biomass observed (Figure 4B) is through an expression-independent effect of Sir2.

**FIGURE 3.**
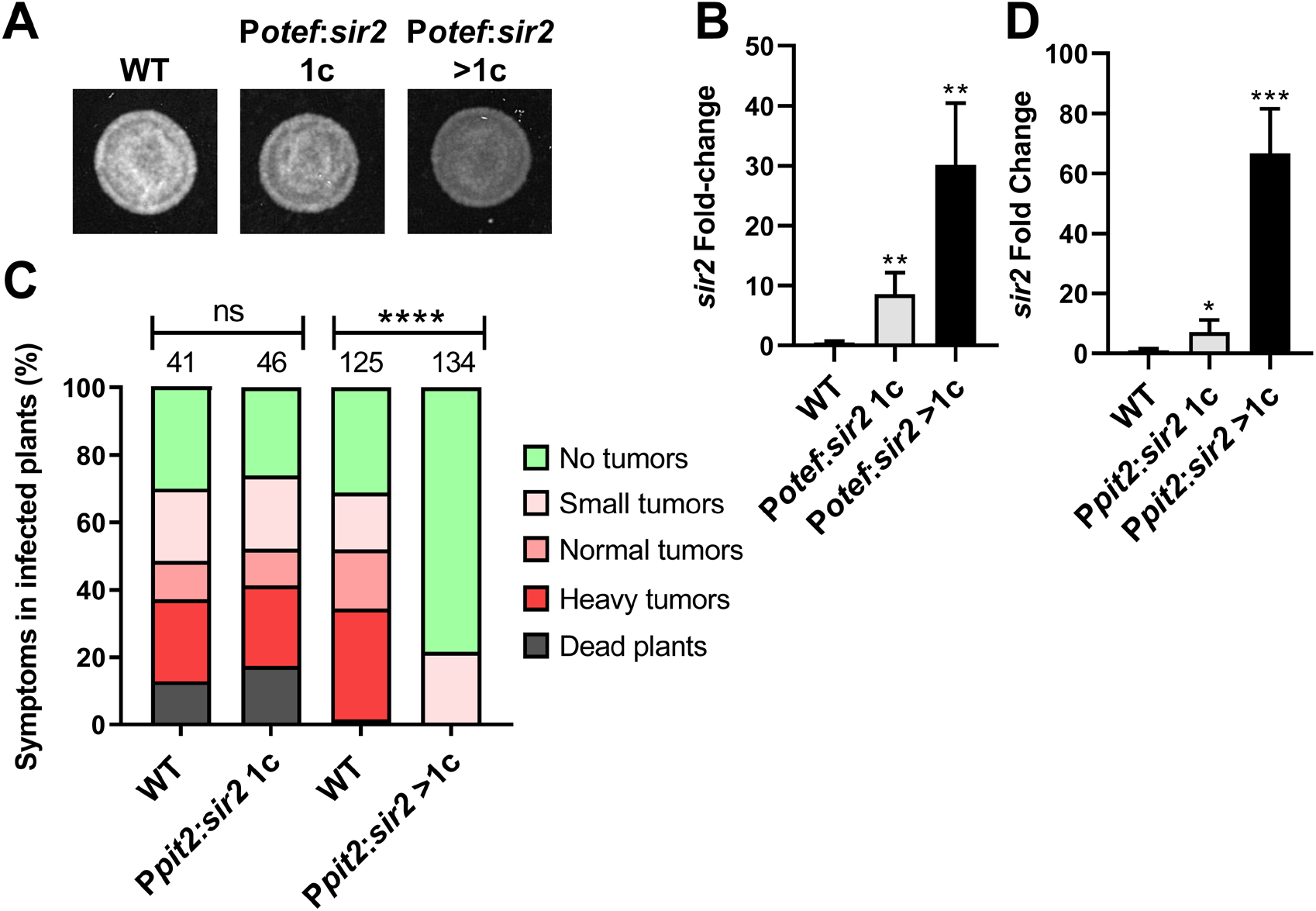
Sir2 overexpression reduces filamentation and virulence in *Ustilago maydis*. **(A)** Filamentation of wild-type and *sir2* overexpression mutants containing one (1c) or more copies (>1c) of the P*otef*:*sir2* construct, grown on PD–charcoal plates for 20 hours at 25ºC. **(B)** *sir2* expression levels in axenic culture of wild-type and the *sir2* overexpression mutants measured by RT-qPCR. *U. maydis ppi1* was used as reference gene. Values were normalized to wild-type. Error bars represent the standard deviation from at least three independent replicates. Student’s t-test statistical analysis was performed (** p-value < 0.005). **(C)** Quantification of symptoms for plants infected with the indicated strains at 14 dpi. Total number of infected plants is indicated above each column. Three biological replicates were analyzed. Mann– Whitney statistical test was performed (ns, no significant; **** p-value < 0.001). **(D)** *sir2* expression levels of wild-type and the *sir2* overexpression mutants containing one (1c) or more copies (>1c) of the P*pit2*:*sir2* construct infecting maize leaves at 3 dpi, measured by RT-qPCR. *U. maydis ppi1* was used as reference gene. Values were normalized to wild-type. Error bars represent the standard deviation from at least three independent replicates. Student’s t-test statistical analysis was performed (** p-value < 0.005).

**FIGURE 4.**
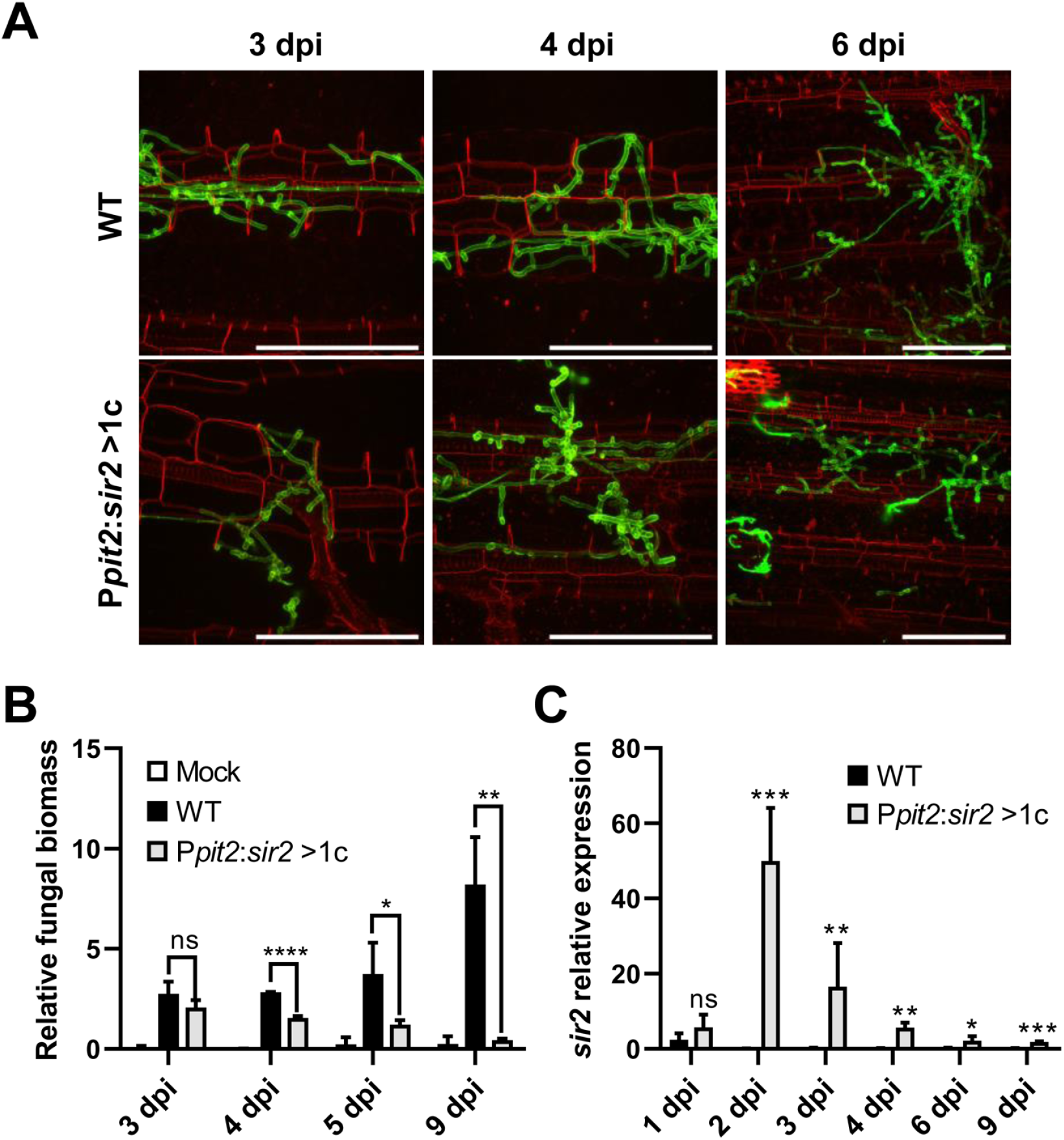
Progression inside the plant is impaired by the overexpression of *sir2*. **(A)** Maize leaves from plants infected with wild-type and the P*pit2*:*sir2* >1c mutant at 3, 4 and 6 dpi were stained with propidium iodide (red) and *U. maydis* hyphae with WGA-AF-488 (green) and visualized by fluorescence microscopy. Scale bar represents 100 μm. **(B)** Relative fungal biomass was calculated by comparison between *U. maydis ppi1* gene and *Z. mays* glyceraldehyde 3-phosphate dehydrogenase gene (GAPDH), measured by RT-qPCR of genomic DNA extracted from leaves infected with wild-type and P*pit2*:*sir2* >1c mutant at 3, 4, 5 and 9 dpi. Error bars represent the standard deviation from three independent replicates. Student’s t-test statistical analysis was performed (ns, not significant, * p-value <0.05, ** p-value < 0.005, **** p-value < 0.0001). **(C)** *sir2* expression of *U. maydis* wild-type and P*pit2*:*sir2* >1c infecting maize leaves at 1, 2, 3, 4, 6 and 9 dpi, measured by RT-qPCR. *U. maydis ppi1* was used as reference gene. Values were normalized to wild-type. Error bars represent the standard deviation from at least three independent replicates. Student’s t-test statistical analysis was performed (ns, not significant, * p-value <0.05, ** p-value < 0.005, *** p-value < 0.0005).

### Sir2 prevents induction of a pool of virulence genes

To study the effect of Sir2 overexpression on gene transcription during infection, we carried out RNA-seq analysis of maize leaves infected with the wild-type strain or the mutant with more than one copy of P*pit2*:*sir2* at 3 dpi, when there was no significant change in fungal biomass and *sir2* had been highly induced. We identified 51 genes downregulated and 39 upregulated in the mutant compared to the wild-type strain (Figure 5A and Supplementary Table S7), with the downregulated genes showing a stronger change in terms of differential expression. When we considered the distribution of transcriptional fold changes in a wild-type strain at 3 days post-infection in comparison to axenic conditions, the downregulated genes in the *sir2*-overexpressed mutant represented a small group of all genes induced during infection (Figure 5B, Supplementary Table S7 and Supplementary Table S8).

**FIGURE 5.**
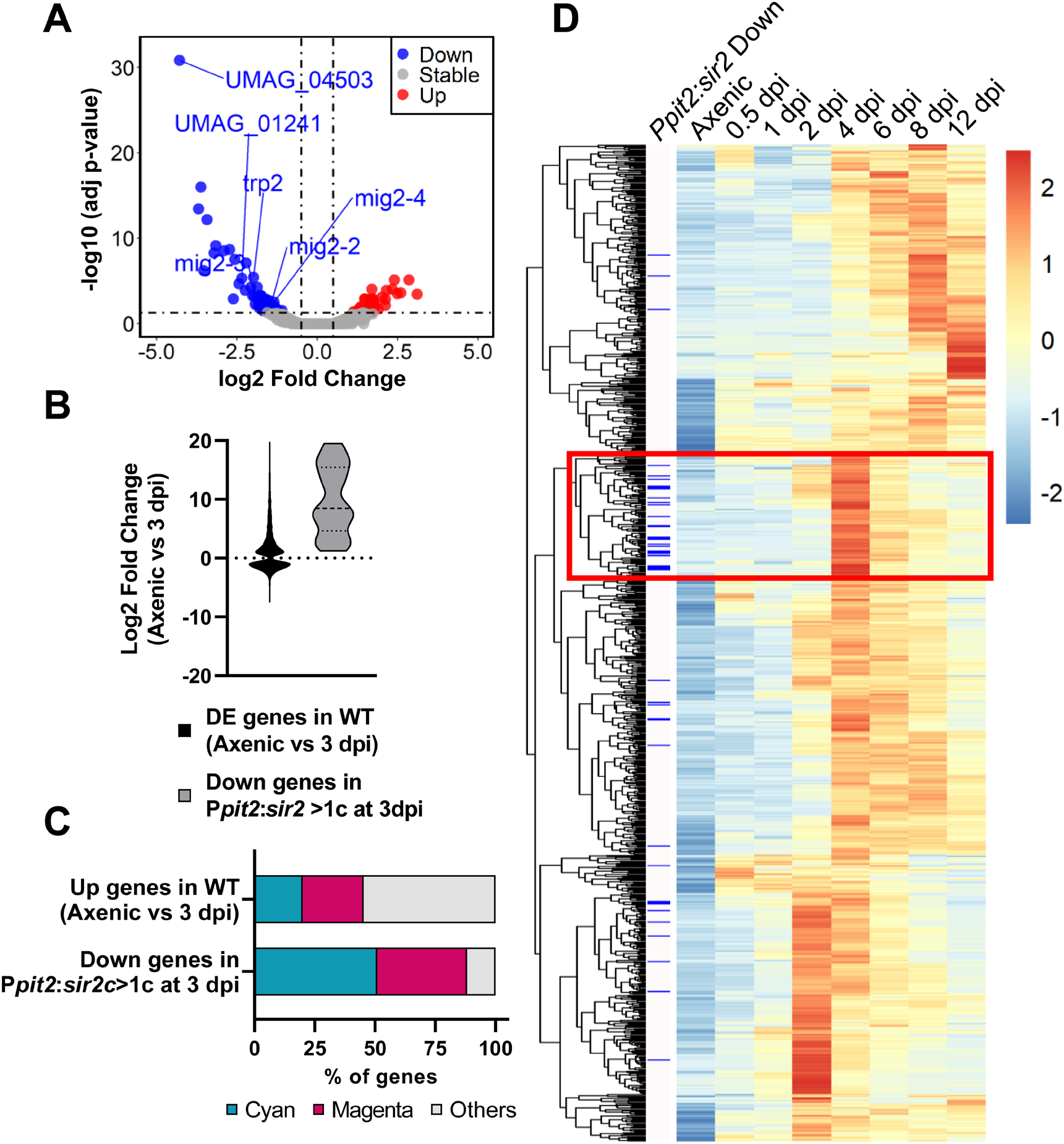
Overexpression of *sir2* during pathogenesis avoids the proper activation of a subpopulation of genes related to biotrophism establishment and tumorigenesis. **(A)** Volcano plot showing the log2 fold change in gene expression and the statistical significance of the differential expression analysis from RNA-seq data obtained in the P*pit2*:*sir2* >1c mutant compared to wild-type at 3 dpi. Red, blue and grey dots represent the upregulated genes (log2 fold change ≥ 0.5, adj p-value < 0.05), downregulated genes (log2 fold change ≤ −0.5 adj p-value < 0.05) and genes without changes in the P*pit2*:*sir2* >1c mutant, respectively. *sir2* and *sdh2* (integration resistance cassette used for over expression mutant) data have been removed for plotting purpose. **(B)** Log2 fold change distribution of the differentially expressed genes (adj p-value < 0.05) at 3 dpi compared to axenic conditions in wild-type (green) and the downregulated genes in the P*pit2*:*sir2* >1c mutant compared to wild-type at 3dpi (blue). **(C)** Percentage of genes belonging to the magenta or cyan modules of coexpressed genes during infection for the upregulated genes of wild-type at 3 dpi compared to axenic culture and for the downregulated genes in the P*pit2*:*sir2* >1c mutant compared to wild-type at 3 dpi. **(D)** Clustering analysis of the expression profile of genes belonging to the cyan and magenta modules in axenic cultures and the indicated dpi for the wild-type strain (normalized counts obtained from the published RNA-seq data (Lanver et al., 2018). Heatmap color-scale values correspond to the Z-score transformation of the expression data, with +2 (red) being the highest expression and -2 (blue) being the lowest expression. Blue bars on the left indicate the downregulated genes in the P*pit2*:*sir2* >1c mutant at 3 dpi compared to wild-type.

Interestingly, as observed in filamentation conditions (Figure 2F), we detected strong enrichment for genes belonging to the cyan (50%) and the magenta (40%) modules in this group of downregulated genes (Figure 5C, Supplementary Table S7 and Supplementary Table S8). Although Sir2 avoided the full activation of mainly cyan and magenta genes, they were only a small subgroup of the entire modules (19 out of 558 magenta genes showed upregulation in our experiment at 3 dpi, and 26 out of 444 total cyan genes) (Supplementary Table S7 and Supplementary Table S8). We wished to determine whether this group of genes has some specific expression profile during infection; therefore, we performed clustering analysis using the expression level of the cyan and magenta gene modules obtained from the RNA-seq data from Lanver *et al*., 2018 (Lanver et al., 2018). In the resulting heatmap, we marked the Sir2-repressed genes, observing that many of them were clustered together (Figure 5D), indicating that they share a similar expression profile. Specifically, they are genes repressed during the first few days of infection and are strongly induced at 4 dpi, several days after *sir2* overexpression. These data may suggest that the overexpression of *sir2* avoids the subsequent induction of a group of genes induced at 4 dpi, which could explain the decrease in fungal biomass that we observed from this day onwards (Figure 4B). However, it is necessary to exercise caution regarding our timing interpretation, as the data obtained by Lanver *et al*., 2018 (Lanver et al., 2018) were obtained in a different genetic background (FB1xFB2), and the timing of the infection process may not be the same as the one that we observed in an SG200 background, where we conducted the *sir2* overexpression.

### Δ*sir2* mutant shows increased acetylation of histone H4 at regulated genes

In order to check whether Sir2 controls filamentation and gene expression through histone deacetylation, we performed Western blot analysis using antibodies against acetylated histone 3 (H3ac) and histone 4 (H4ac), the canonical histone targets of Sir2 (Robyr et al., 2002; Suka et al., 2002; Vaquero et al., 2006; Shimizu et al., 2012; Cai et al., 2021; Zhao and Rusche, 2022), from total proteins extracted after growth in rich media (Figure 6A) and in filamentation induction media (Figure 6B). We did not detect any significant change for these two modifications in either a *sir2* deletion or overexpression mutant (Figures 6A, B). Due to the effect observed in the *sir2* mutants for specific loci rather than broad chromatin regions, we studied the acetylation state of different Sir2 regulated genes. To this aim, we carried out Chromatin Immunoprecipitation (ChIP) experiments using antibodies against H3ac and H4ac followed by RT-qPCR of the promoter region and the ORF of some of the genes that change their expression in the Δ*sir2* mutant in axenic culture (*eff1-9*) (Figure 2C, Supplementary Figure S2.C and Supplementary Table S4), PD-charcoal plates (UMAG_06128, *mig2-6, rsp3*) (Figure 2D and Supplementary Table S5) and at 3 dpi in the *sir2* overexpressed mutant (*mig2-3*, UMAG_01241) (Figure 5A and Supplementary Table S7). As observed in Figure 6C, there was an enrichment of acetylated H4 in *eff1-9* and UMAG_06128 and a slight increase of acetylated H3 in the promoter region of *eff1-9* in the Δ*sir2* mutant. As Sir2 commonly deacetylase lysine 16 in histone 4 (H4K16) (Robyr et al., 2002; Suka et al., 2002; Vaquero et al., 2006; Shimizu et al., 2012; Cai et al., 2021; Zhao and Rusche, 2022), we checked the H4K16 acetylation level by ChIP and RT-qPCR of the genes with enriched H4 acetylation, but no differences were observed in the acetylation of this residue (Figure 6D).. Because histone acetylation has been described to be dependent on transcription level (Martin et al., 2021), we checked the reads of the selected target genes obtained in our RNA-seq analysis in axenic condition and observed a correlation between the upregulation of expression on the Δ*sir2* mutant and the increase in acetylation (Figures 6C, E). These data suggest that the increase in histone acetylation is a consequence and not the cause of the gene upregulation and that the defects observed in filamentation and gene regulation in this work are due to a deacetylation-independent function of Sir2 or the deacetylation of a different target.

**FIGURE 6.**
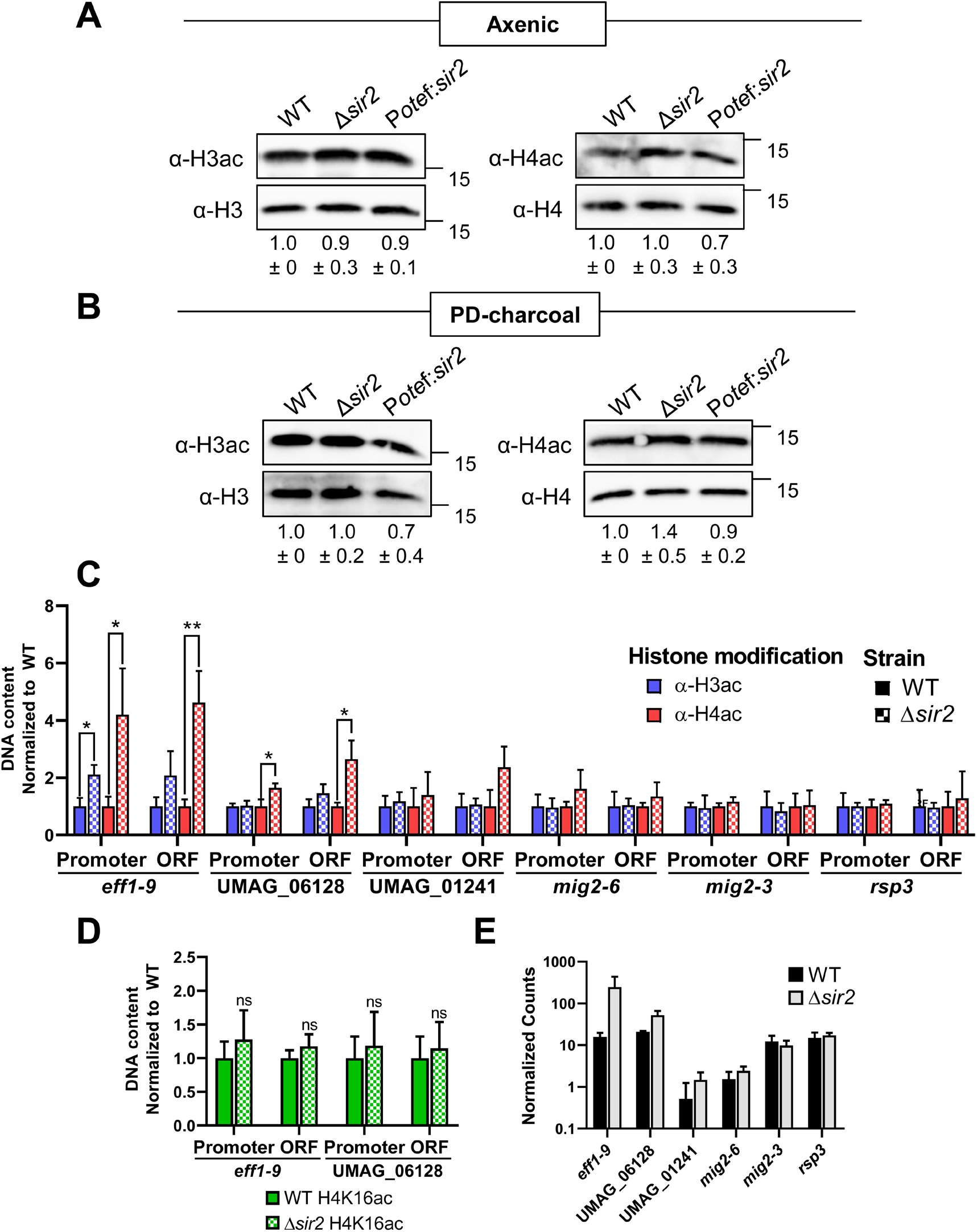
Effect of *sir2* mutants in histone acetylation. **(A, B)** Total proteins extracted from axenic culture **(A)** or PD-charcoal plates **(B)** of wild-type, Δ*sir2* and the P*otef*:*sir2* >1c strains were used for Western blotting. The H3ac and H4ac antibodies were used to detect histone acetylation and H3 and H4 antibodies were used as loading controls. H3ac and H4ac signals were normalized to H3 and H4 levels, respectively, and compared to wild-type from three independent replicates. **(C, D)** ChIP analysis using H3ac and H4ac antibodies **(C)** or H4K16ac antibody **(D)** on chromatin extracts from axenic culture of wild-type and Δ*sir2* strains. Immunoprecipitated DNA was analyzed by RT-qPCR, amplifying regions within the promoter and the open reading frame (ORF) of the indicated gene. Values correspond to the amount of DNA recovered in the IP relative to *ppi1* gene. Values were normalized to wild-type. Error bars represent the standard deviation from at least three independent replicates. Student’s t-test statistical analysis was performed (ns, no significant, * p-value <0.05, ** p-value < 0.005). **(E)** Mean of the normalized count of the indicated genes obtained in the RNA-seq experiment in the wild-type and Δ*sir2* strains in axenic condition.

## Discussion

Sirtuins are NAD^+^-dependent deacetylases with important regulatory roles in processes such as lifespan, metabolic control or pathogenesis in fungi. As they require the NAD+ cofactor for their activity, sirtuins may serve as regulators of many of these processes in response to metabolic stages. Here, we have performed a systematic analysis of all sirtuins present in *U. maydis* and focused on the nuclear sirtuin Sir2, demonstrating its role in the control of part of the pathogenic program.

### The sirtuin repertoire of *U. maydis*

Fungal sirtuins can be classified into five principal subfamilies according to their orthologs in other organisms: Sir2/Hst1, Hst2, Hst3/4, SirT4 and SirT5 (Zhao and Rusche, 2022). Interestingly, *U. maydis* harbors a sirtuin member for each subfamily (Elías-Villalobos et al., 2019). The SirT4 and SirT5 subfamilies are mitochondrial sirtuins with not many examples described in fungi. We have demonstrated here that the two members of these families in *U. maydis*, Hst5 and Hst6, display mitochondrial localization. However, we have not detected any essential role for them during pathogenesis. On the other hand, Hst2 has been described to show a primary cytoplasmatic localization, although it is involved in locus-specific silencing (Halme et al., 2004; Durand-Dubief et al., 2007) and has a prominent function in chromosome condensation during mitosis (Vaquero et al., 2006; Wilkins et al., 2014; Kruitwagen et al., 2018; Jain et al., 2021). This conflict between location and function may be explained in mammalian cells since Hst2 can move from the cytoplasm to the nucleus, mainly in premitotic cells (Vaquero et al., 2006; Wilson et al., 2006). Similarly to mammalian cells, we have observed that *U. maydis* Hst2 is mainly cytoplasmic, but it is transported to the nucleus in premitotic cells and it remains bound to chromatin during mitosis. This observation strongly supports the idea that in *U. maydis*, Hst2 conserves its role in chromatin condensation during mitosis, probably via the deacetylation of H4K16, as in yeast and mammals (Vaquero et al., 2006; Wilkins et al., 2014). Finally, we found Hst4 and Sir2, members of the Hst3/4 and Sir2/Hst1 subfamilies, respectively, both with constitutive nuclear localization. Although Hst4, as well as the other member of the subfamily, Hst3, may be involved in the transcriptional silencing of a specific locus and heterochromatin (Freeman-Cook et al., 1999; Durand-Dubief et al., 2007), the main function described is their role in chromosome integrity through H3K56 deacetylation. Consequently, cells with a misregulation of these sirtuins show genome stability-related phenotypes such as spontaneous DNA damage, chromosome loss or sensitivity to DNA damage (Celic et al., 2006; Maas et al., 2006; Haldar and Kamakaka, 2008). Although mutants for *hst3* and *hst4* are viable in *S. cerevisiae* and *S. pombe*, the deletion of the single member of this subfamily in *Candida albicans, hst3*, which has been demonstrated to be involved in H3K56 deacetylation, has not been possible (Wurtele et al., 2010), suggesting that this gene is essential in this yeast. Here, we have demonstrated that the deletion of *hst4* in *U. maydis* is lethal, which supports again a conserved role of Hst4 in this fungus in genome stability through the acetylation of H3K56. The other nuclear sirtuin in *U. maydis* is Sir2. This is the founding member of the family and has been extensively characterized in different organisms. The main described role for this sirtuin is transcriptional silencing through histone deacetylation, particularly of H4K16 and H3K9 (Robyr et al., 2002; Suka et al., 2002; Vaquero et al., 2006; Shimizu et al., 2012; Cai et al., 2021; Zhao and Rusche, 2022). The most recognized biological function for this sirtuin is the regulation of aging (Kaeberlein et al., 1999; Fabrizio et al., 2005; Fu et al., 2008; Bouklas et al., 2017). However, in the human pathogens *Candida glabrata* (Domergue et al., 2005) and *Cryptococcus neoformans* (Arras et al., 2017), in the insect pathogen *Beauveria bassiana* (Cai et al., 2021) and in the plant pathogen *Magnaporthe oryzae* (Fernandez et al., 2014), it has been demonstrated to have implications for the virulence process. In *M. oryzae*, thus far, the only phytopathogen with an in-depth characterization of Sir2, this sirtuin controls virulence through deacetylation of the transcriptional repressor Jmjc, which allows the expression of the superoxide dismutase Sod1, with important roles in ROS detoxification during the first few steps of infection (Fernandez et al., 2014). Here, we provide another example of the role of Sir2 in a phytopathogen, now a basidiomycete, where this sirtuin affects the regulation of a subset of the virulence genes. We have observed that the overexpression of *sir2* reduces the filamentation capability and inhibits the correct induction of virulence genes, affecting proper tumor formation. In contrast, the deletion of *sir2* causes an increase in the filamentation and infection capabilities and allows a better induction of filamentation and virulence genes. Interestingly, Sir2 affects filamentation as well in the human pathogen *C. albicans* (Zhao and Rusche, 2021). However, the effect is the reverse, as Sir2 is essential for proper filamentation in this yeast. Further examples of the role of Sir2 in filamentation in different fungi would be useful to verify the possible conserved role of Sir2 during this process.

### The role of UmSir2 in transcriptional regulation

Sir2 has a transcriptional silencing role conserved through evolution (Hickman et al., 2011). In high eukaryotes and yeast such as *S. pombe*, with the hallmarks of high eukaryotes and heterochromatin (methylation of H3K9, Heterochromatin Protein 1 (HP1) and RNA interference (RNAi) to produce the heterochromatin silencing platform), Sir2 has been described to aid in the silencing of this region through the deacetylation of histones, mainly H4K16 and H3K9 (Shankaranarayana et al., 2003). However, *U. maydis* lacks all these hallmarks of heterochromatin formation. A similar situation is found in *S. cerevisiae*, where RNAi, H3K9 methylation and HP1 proteins are not present (Hickman et al., 2011). In this organism, Sir2, as part of the SIR complex, together with Sir3 and 4, is the main factor involved in the formation of heterochromatin-like regions (Robyr et al., 2002; Suka et al., 2002). It is tempting to believe that in *U. maydis*, with a similar chromatin scenario, Sir2 would have a similar central role in the silencing of heterochromatin-like regions. However, in *U. maydis*, we did not find homologs of the other members of the SIR complex, which are required for the spreading effect required for heterochromatin formation (Elías-Villalobos et al., 2019). In addition, in a mass spectrometry analysis of a Sir2:eGFP pull down (Supplementary Figure S4), we have not detected any factor belonging to a putative SIR complex. Thus, we observe a different scenario in *U. maydis*, with no hallmarks of heterochromatin and no apparent SIR complex. Our RNA-seq studies reveal that Sir2 seems to not be an essential factor for heterochromatin-like silencing in this organism, as we have not found a clear effect in the characteristic heterochromatin regions, such as telomeres or centromeres, or in clusters of coregulated genes, which would be putative chromatin-silenced areas. However, we have found a direct or indirect regulatory effect of Sir2, mainly in silencing, in specific loci. This is consistent with the regulatory effect observed for different sirtuins, including Sir2, and in the general silencing effect observed through evolution (Zhao and Rusche, 2022).

An interesting observation is that Sir2 is degraded during filamentation, and we were able to detect two bands of this protein by Western blot (Figure 2A). A tentative speculation is that a posttranslational modification (ptm) of Sir2 during the pathogenic program leads to its degradation and/or inactivation, allowing the proper expression of a pool of virulence genes. Sir2 has been described in yeast to suffer phosphorylation and sumoylation, altering its function (Hannan et al., 2015; Kang et al., 2015). In *M. oryzae*, Sir2 accumulation during infection is controlled through ubiquitination by the E3 ubiquitin ligase Upl3, and basal Sir2 levels are controlled by the Grr1 (Patton et al., 1998) and Ptr1 ones (Li et al., 2020). Interestingly, we found UMAG_04611, a homolog of the E3 ubiquitin ligase Skp1, to interact with Sir2 in our mass spectrometry analysis. Further analysis would be interesting to confirm the biological significance of this interaction and the possible role of ubiquitination or other Sir2 modifications in *U. maydis* filamentation and virulence.

### Possible substrates of Sir2

As mentioned previously, Sir2 represses the transcription of specific substrates or mediates the silencing of heterochromatin regions by the deacetylation of histones, typically H4K16 and H3K9 (25– 27, 30, 63, 64). In *U. maydis*, in agreement with the observed repressive effect in specific loci for Sir2, we have not detected any significant change in acetylation for either H3 or H4 by Western blot. However, there is a significant increase of mainly H4 acetylation when specific regulation targets of Sir2 were studied by ChIP and RT-qPCR. Interestingly, we have not observed an increase in acetylation for the specific lysine 16. Although we cannot discard the possibility of a different lysine residue as the target for Sir2, the correlation between the increase in acetylation and the transcriptional upregulation observed, which may indicate an indirect effect in H4 acetylation due to the increase in transcription, suggests a different protein as the target of Sir2 in *U. maydis*. In agreement with this, in the insect pathogen *B. bassiana*, acetylation analysis has revealed that, besides H4 and H3, hundreds of proteins show an altered acetylation pattern in a *sir2* mutant (Cai et al., 2021). More specifically, in plant pathogens, Sir2 has been demonstrated to affect the virulence of *M. orizae* by deacetylation of a transcription factor (Fernandez et al., 2014). In our mass spectrometry analysis, we found the transcription factor Med1 to interact with Sir2 (Supplementary Figure S4). This transcription factor acts upstream of the virulence master regulator Prf1, affecting its expression and virulence (Chacko and Gold, 2012). Although we have not found any significant defect in *prf1* expression on a *sir2* mutant, further investigation would be necessary to study a possible regulatory effect of Sir2 over the Med1 transcription factor and other possible substrates that could explain more in depth the filamentation and transcriptional defects observed on the *sir2* mutant in *U. maydis*.

## Author Contributions

RRB and JII designed the concept of the study and supervised the project. BN performed the experiment. RRB and BN analyzed the data. RRB and BN drafted the manuscript. All authors contributed to the article.

## Acknowledgements

We thank the Genetics Department for their useful discussions and comments; Víctor Manuel Carranco, Sandra Romero, Laura Tomás and Ismael Fernández Portillo for technical support.

## Data availability

RNA sequencing data has been submitted to NCBI Genbank and are available under the following links: httpsXXXXXXXXX

