## Supplemental Figures for "Systematic characterization of *Ustilago maydis* sirtuins shows Sir2 as a modulator of pathogenic gene expression"

### Supplementary Figure S1

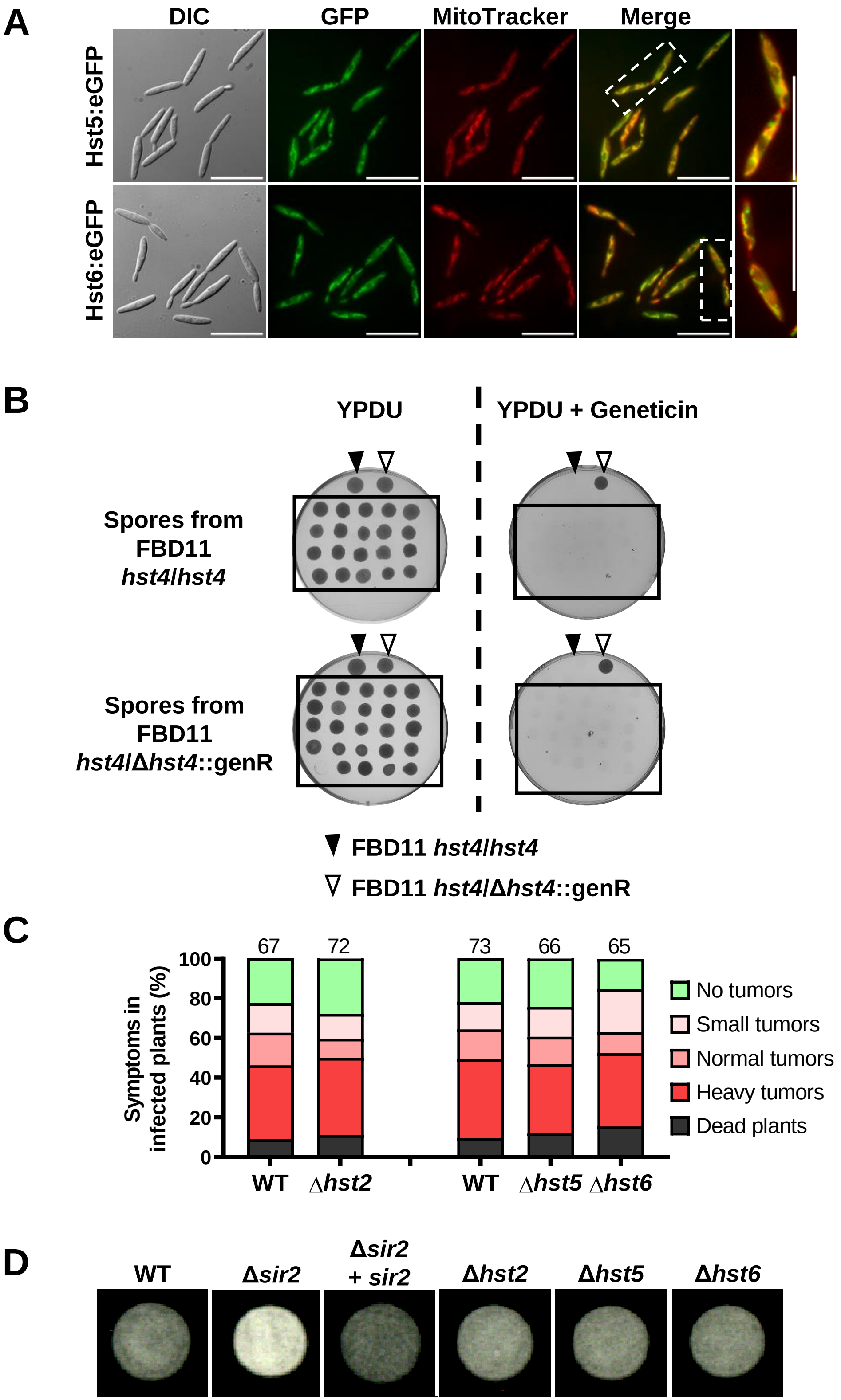

**SUPPLEMENTARY FIGURE S1**. Hst5 and Hst6 localize in the mitochondria, deletion of *hst4* is not viable for *Ustilago maydis* and Hst2, Hst5 and Hst6 have no effects in the virulence of the fungus. **(A)** Confocal microscopy of *U. maydis* strains expressing the indicated sirtuins tagged with eGFP in its own loci. Mitochondria were stained with MitoTracker. Scale bar represents 20 µm. **(B)** FBD11 wild-type and FBD11 *hst4*/Δ*hst4*::genR diploid strains were inoculated on maize plant. None of the singularized colonies obtained from germinated spores were able to grow on selective medium (YPDU plus geneticin). YPDU plates were used as control. Control strains were spotted on top of the plates. **(C)** Quantification of symptoms for plants infected with the indicated strains at 14 dpi. Total number of infected plants is indicated above each column. Two biological replicates were analyzed. Mann–Whitney statistical test was performed (ns, no significant; **** p-value < 0.001). **(D)** Filamentation of wild-type and the indicated sirtuins mutants grown on PD-charcoal plates for 18 hours at 25ºC.

### Supplementary Figure S2

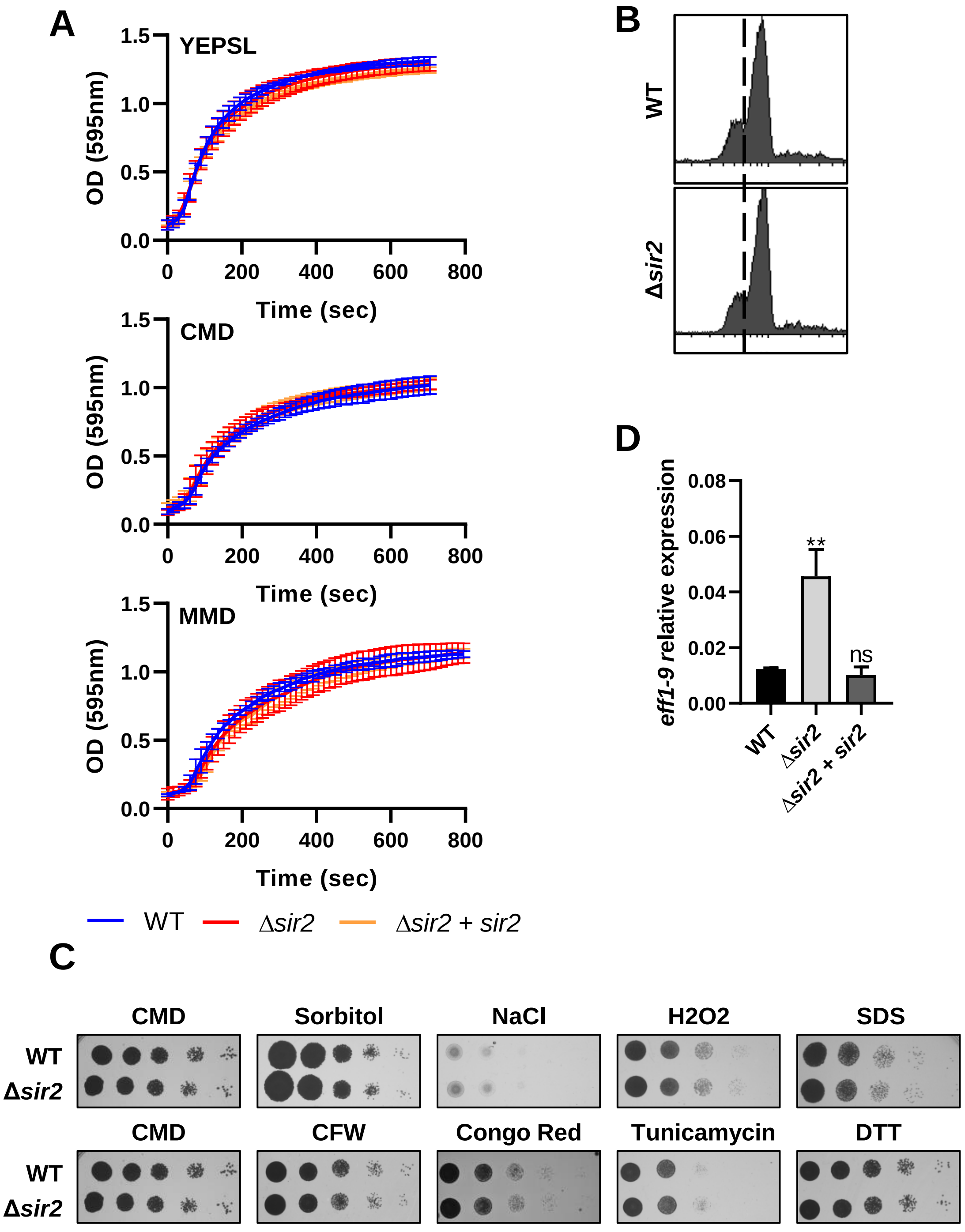

**SUPPLEMENTARY FIGURE S2**. Deletion of *sir2* has not pleiotropic effects and is complemented by reintroducing *sir2* gene. **(A)** Growth curve of wild-type and Δ*sir2* mutant growing in YEPSL, CMD or MMD media. Error bars represent the standard deviation from three independent replicates. **(B)** Flow cytometry analysis of the DNA content of wild-type and Δ*sir2* mutant grown in CMD medium. Relative fluorescence intensities are given on horizontal axes and vertical axes reflect cell numbers. **(C)** Spot tests to assay osmotic stress (sorbitol and NaCl), oxidative stress (H2O2), membrane integrity (SDS), cell wall integrity (calcofluor white (CFW) and Congo red) and endoplasmic reticulum stress (Tunicamycine and DTT). CMD without drug was used as growth control. **(D)** *eff1-9* expression in axenic culture of wild-type, Δ*sir2* and the Δ*sir2* complementation strains measured by RT-qPCR. U. maydis *ppi1* gene was used for normalization. Error bars represent the standard deviation from three independent replicates. Student's t-test statistical analysis was performed (ns, not significant; ** p-value < 0.005).

### Supplementary Figure S3

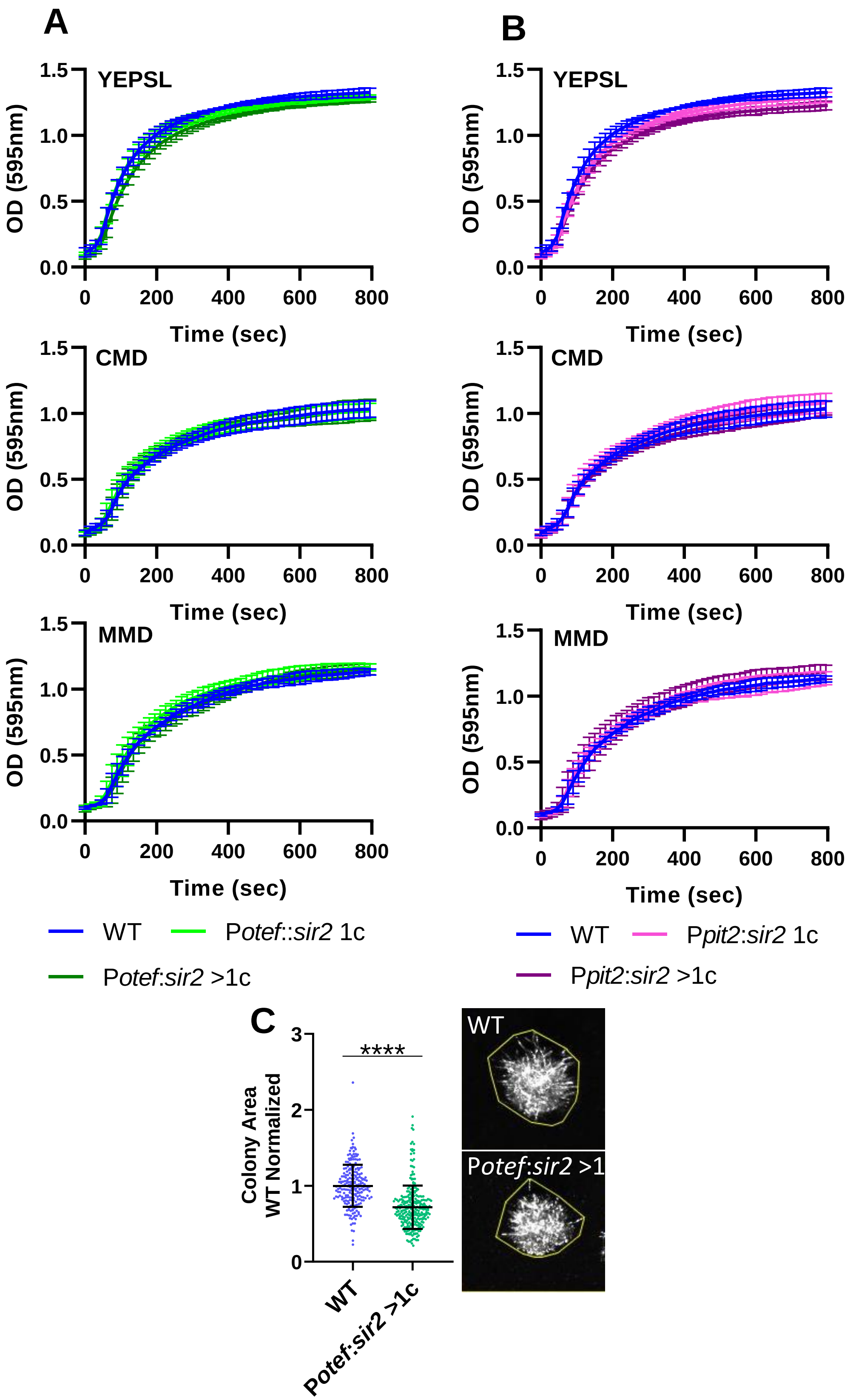

**SUPPLEMENTARY FIGURE S3**. Sir2 overexpression does not affect cell growth but reduces the filament formation on PD-charcoal plates. **(A, B)** Growth curve of wild-type, Potef:sir2 1c and Potef:sir2 >1c mutants **(A)** or Ppit2:sir2 1c and Ppit2:sir2 >1c mutants **(B)** growing in YEPSL, CMD or MMD media. Error bars represent the standard deviation from three independent replicates. **(C)** Quantification of the area of the wild-type and Δ*sir2* mutant single colonies grown on PD–charcoal plates for 48 hours at 25ºC. The colony area was measured as indicated in the stereoscopic images. Data was normalized with the mean of the area of the wild-type colonies. Three biological replicates were analyzed. Student's t-test statistical analysis was performed (**** p-value < 0.001).

### Supplementary Figure S4

A

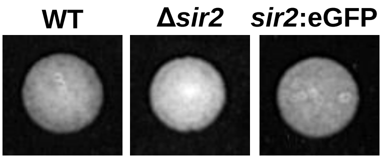


B

| GeneID | Gene Name | Description | Peptides in Sir2:eGFP | Peptides in wild-type |
|---|---|---|---|---|
| UMAG_02845 | vps54 | Vacuolar protein sorting-associated protein 54 | 3 | 0 |
| UMAG_11692 | arp1 | Centractin | 3 | 0 |
| UMAG_10587 | ppx1 | Exopolyphosphatase | 3 | 0 |
| UMAG_03588 | med1 | Transcription factor medusa | 2 | 0 |
| UMAG_04611 | skp1 | E3 ubiquitin ligase complex SCF subunit | 2 | 0 |
| UMAG_05101 | top1 | DNA topoisomerase I | 2 | 0 |
| UMAG_01122 | hcs1 | DNA helicase A | 2 | 0 |
| UMAG_10855 | mkk1 | MAPK signalling pathway | 2 | 0 |
| UMAG_02883 | pfdn1 | Prefoldin subunit 1 | 2 | 0 |
| UMAG_00189 | esf1 | 18S rRNA factor | 2 | 0 |
| UMAG_05552 | | Hypothetical protein | 2 | 0 |
| UMAG_11756 | | Hypothetical protein | 2 | 0 |
| UMAG_03419 | | Hypothetical protein | 2 | 0 |
| UMAG_10436 | | Hypothetical protein | 2 | 0 |
| UMAG_03789 | | Hypothetical protein | 2 | 0 |

**SUPPLEMENTARY FIGURE S4.** Sir2 interacting proteins. **(A)** Sir2 fussed to eGFP does not affect its function. Images showing the filamentation of wild-type, and the indicated mutants on PD-charcoal plates for 18 hours at 25ºC. **(B)** Proteins interacting with Sir2. Identified proteins by Sir2 immunoprecipitation followed by mass spectrometry analysis. Proteins with log2 >6 and with, at least, 2 peptide in all replicates of the *sir2*:eGFP mutant and none in the wild-type are shown.

### Supplementary Methods

#### Supplementary Method S1. Growth curve and stress assay.

For the growth curve assay, U. maydis cells were grown to the exponential phase and diluted to OD600 of 0.1 in YEPSL, CM supplemented with 1% D-glucose (CMD) or MM supplemented with 1% D-glucose (MMD) (Holliday, 1974). Cell growth at 28ºC with continuous shaking was analyzed over 24 h using a Spark 10M fluorescence microplate reader (Tecan, Männedorf, Switzerland). Cell wall integrity, membrane integrity, osmotic, endoplasmic reticulum and oxidative stress assays were carried out with cultures grown at 28º C to the exponential phase in CMD and spotted at 0.4 OD600 onto CMD plates supplemented with calcofluor white (CFW) 40 µg/mL (Sigma-Aldrich, Darmstadt, Germany), Congo Red 50 µg/mL (Sigma-Aldrich, St. Louis, MO, USA), 4 mM DTT (iNtRON Biotechnology, Seongnam, Gyeonggi, ROK), tunicamycin 1 µg/mL (Sigma-Aldrich, Darmstadt, Germany), sorbitol 1 M (Sigma-Aldrich, Darmstadt, Germany), 2% DMSO (Sigma-Aldrich, Darmstadt, Germany), H2O2 0.75 mM (Sigma-Aldrich, Darmstadt, Germany), NaCl 1 M (Sigma-Aldrich, Darmstadt, Germany) and 0.005% SDS (Sigma-Aldrich, Darmstadt, Germany). Plates were incubated at 28º C for 48 h.

#### Supplementary Method S2. Spore germination.

Spore germination was conducted according to published protocols (Eichhorn et al., 2006). Mature tumors of plants infected with U. maydis FBD11 strain and it derivate FBD11 Δhst4 were harvested 21 dpi and dried at 37ºC for 2 days. Spores were rehydrated with distilled water, crushed with a mortar and incubate with 3% CuSO4 solution at RT for 4 hours. Washed spores were germinated in YPDU plated supplemented with Ampicillin (100 µg/ml), Chloramphenicol (25 µg/mL) and Tetracycline (10 µg/ml). To test FBD11 Δhst4 viability, singularized colonies from germinated spores were grown in YPDU supplemented with geneticin (2 µg/ml).

#### Supplementary Method S3. Flow cytometry.

To analyze the Δsir2 mutant DNA content, we followed the previously described protocol (García-Muse et al., 2003). Cells were grown in CMD medium to exponential phase. Prior to analysis, cells were harvested, washed twice with cold water, fixed in 70% ethanol overnight, and resuspended in 50 mM sodium citrate, pH 7.5. Cellular RNA was eliminated by incubation with RNase A (0.25 mg/mL) at 50ºC for 1 hour and then cells were incubated with proteinase K (1 mg/mL) for 1 hour at 50ºC. Cells were stained with propidium iodide (16 mg/mL) and the fluorescence of 10.000 cells was measured using a FACSCalibur flow cytometer (Becton Dickinson, East Rutherford, NJ, USA) with a 530/30 bandpass filter.

#### Supplementary Method S4. Immunoprecipitation and mass spectrometry.

Cells grown in YEPSL to exponential phase were collected and washed twice with 20 mM Tris-HCl pH 8.8. Pellets were then resuspended, and cells lysed in RIPA buffer (50 mM Tris/HCl, pH 8, 150 mM NaCl, 1% Nonidet P-40, 0.5% sodium deoxycholate, 0.1% SDS) supplemented 1 µg/ml Pepstatin A (PanReac AppliChem, Barcelona, Spain), 1 μg/ml Bestatin (Thermo Scientific, Carlsbad, CA, USA), 1mM PMSF (PanReac AppliChem, Barcelona, Spain) and EDTA-free protease inhibitor complex (cOmplete Tablets EDTA-free, Roche, Mannheim, BW, Germany). After cell lysis, samples were centrifuged at 14000 rpm for 30 min at 4ºC and the supernatant was collected. Sir2:GFP was immunoprecipitated with ChromoTek GFP-Trap® Magnetic Agarose (ChromoTek, Planegg, Germany). Elution was performed by boiling the samples in Laemmli buffer supplemented 200 mM DTT and proteins precipitated by addition of 4 volumes 1:1 methanol:acetone. Protein pellets were resuspended in urea 6M, 50mM ammonium bicarbonate. Disulphide bonds were reduced adding DTT 10Mm and for carbamidometylation of cysteine –SH groups, IAA 30 mM was added. Samples were digested overnight at 37 ºC using trypsin bovine (Sequencing Grade Modified Trypsin, Promega) in a ratio 1:12 enzyme-substrate. Reaction was stopped using formic acid to 0.5%. OMIX C18 tips (Agilent Technologies) were used for concentrating and desalting peptide extracts. Samples were dried and resuspended in 0.1% trifluoroacetic acid. 1ug of each sample was injected in nano-HPLC system. LC-MS data were analyzed using the SEQUEST® HT search engine in Thermo Scientific™ Proteome Discoverer™ 2.2 software. Data were searched against the Uniprot Ustilago maydis protein database and results were filtered using a 1% protein FDR threshold. Two replicates of each strain were processed.

#### Supplementary Method References


Eichhorn, H., Lessing, F., Winterberg, B., Schirawski, J., Kämper, J., Müller, P., et al. (2006). A ferroxidation/permeation iron uptake system is required for virulence in *Ustilago maydis*. *Plant Cell* 18, 3332–3345. doi: 10.1105/tpc.106.043588.

García-Muse, T., Steinberg, G., and Pérez-Martín, J. (2003). Pheromone-induced G2 arrest in the phytopathogenic fungus *Ustilago maydis*. *Eukaryot. Cell* 2, 494–500. doi: 10.1128/EC.2.3.494-500.2003.

Holliday, R. (1974). "*Ustilago maydis*," in *Bacteria, Bacteriophages, and Fungi* (Boston, MA: Springer US), 575–595. doi: 10.1007/978-1-4899-1710-2_31.
