## Supplemental Table S1-3 for "Systematic characterization of *Ustilago maydis* sirtuins shows Sir2 as a modulator of pathogenic gene expression"

**TABLE S1. Strains used in this study.**

| **Strain** | **Relevant genotype** | **Reference** |
| --- | --- | --- |
| SG200 | *a1:mfa2 bW2bE1* | Bölker et al., 1995 |
| SG200 Δ*sir2* | *a1:mfa2 bW2bE1* Δ*sir2*::nat | This work |
| SG200 Δ*hst2* | *a1:mfa2 bW2bE1* Δ*hst2*::nat | This work |
| SG200 Δ*hst5* | *a1:mfa2 bW2bE1* Δ*hst5*::gen | This work |
| SG200 Δ*hst6* | *a1:mfa2 bW2bE1* Δ*hst5*::hyg | This work |
| SG200 *sir2*:eGFP | *a1:mfa2 bW2bE1* *sir2*:eGFP:hyg | This work |
| SG200 *hst2*:eGFP | *a1:mfa2 bW2bE1* *hst2*:eGFP:hyg | This work |
| SG200 *hst4*:eGFP | *a1:mfa2 bW2bE1* *hst4*:eGFP:hyg | This work |
| SG200 *hst5*:eGFP | *a1:mfa2 bW2bE1* *hst5*:eGFP:hyg | This work |
| SG200 *hst6*:eGFP | *a1:mfa2 bW2bE1* *hst6*:eGFP:hyg | This work |
| SG200 Δs*ir2* + *sir2* | *a1:mfa2 bW2bE1* *sir2*::nat::*sir2*:gen | This work |
| SG200 P*otef*:*sir2* | *a1:mfa2 bW2bE1* *ip*R[P*otef*:*sir2*]*ip*S | This work |
| SG200 P*pit2*:*sir2* | *a1*:*mfa2 bW2bE1* *ip*R[P*pit2*:*sir2*]*ip*S | This work |
| FBD11 | a1a2/b2b2 | Banuett and Herskowitz, 1989 |
| FBD11 Δ*hst4* | a1a2/b2b2 *hst4*/Δ*hst4*::gen | This work |

Banuett, F., and Herskowitz, I. (1989). Different a alleles of *Ustilago maydis* are necessary for maintenance of filamentous growth but not for meiosis. *Proc. Natl. Acad. Sci.* 86, 5878–5882. doi: 10.1073/pnas.86.15.5878.

Bölker, M., Böhnert, H. U., Braun, K. H., Görl, J., and Kahmann, R. (1995). Tagging pathogenicity genes in *Ustilago maydis* by restriction enzyme-mediated integration (REMI). *Mol. Gen. Genet. MGG* 248, 547. doi: 10.1007/BF02423450.

**TABLE S2. Primers used in this study.**

| **Gene** | **Use** | **Name** | **Primer sequence (5´-3´)** |
| --- | --- | --- | --- |
| *sir2* | Deletion construct | sir2KO5_fwd | AAAGGTGCTACTCAACCTCGTCG |
|  |  | sir2KO5_rev | CACGGCCTGAGTGGCCGTAGCAATTCCAAGCCGAGACG |
|  |  | sir2KO3_fwd | GTGGCCATCTAGGCCGAGTACGACACCCAATGGAC |
|  |  | sir2KO3_rev | CGACGCAGCATATCCGAAGC |
|  | Deletion verification | sir2CO5_fwd | AGATGATGATGCCGATGCTACC |
|  |  | nat2_rev | TGTACGCATGTAACATTATACTGAAAACCT |
|  |  | nat1_fwd | TGGCTGCTGATCACAGCAAGTCAGATT |
|  |  | sir2CO3_rev | CATACACGAGCAACGGCAACG |
|  |  | sir2KOint_fwd | TGACAAGACCTTCTTTCTCTCC |
|  |  | sir2KOint_rev | AGTGGAGGATTGGCGTCTTGT |
|  | Complementation | sir2KO5_fwd | AAAGGTGCTACTCAACCTCGTCG |
|  |  | sir2EC5_rev | CACGGCCTGAGTGGCCTCTTCAACAGGAAGGGGAGC |
|  |  | sir2EC5_fwd | GTGGCCATCTAGGCCTCATAGATCACCTCTCTCTC |
|  |  | sir2EC3_rev | TCTCGACATCGGTGACAAGG |
|  | Complementation verification | sir2CO5_fwd | AGATGATGATGCCGATGCTACC |
|  |  | gen1_rev | TCTTCTGAGCGGGACTCTGG |
|  |  | gen2_fwd | GTACGGGTACATCGGATCTGC |
|  |  | sir2CO3_rev | CATACACGAGCAACGGCAACG |
|  |  | sir2KOint_fwd | TGACAAGACCTTCTTTCTCTCC |
|  |  | sir2KOint_rev | AGTGGAGGATTGGCGTCTTGT |
|  | eGFP tagging | sir2ORF_fwd | TCGAGGTCGACGGTATCGATAAGCTTGATAAAGACATGTTTGACAAGACC |
|  |  | sir2ORF_rev | GGTGAACAGCTCCTCGCCCTTGCTCACCATTGATGGAGCCTGTTGTGAATGC |
|  |  | eGFPHyg_sir2_fwd | ATGGTGAGCAAGGGCGAGGA |
|  |  | eGFPHyg_sir2_rev | ATAGGGCGAATTGGAGCTCG |
|  |  | sir2Ter_fwd | CCTGAGTGGCCGAGCTCCAATTCGCCCTATCATCTCCTTCGAGTACGACA |
|  |  | sir2Ter_rev | GCTCTAGAACTAGTGGATCCCCCGGGCTGCACAGCTTCATCCCGTACTGG |
|  |  | sir2GFP_fwd | AAGACATGTTTGACAAGACC |
|  |  | sir2GFP_rev | TGATGGAGCCTGTTGTGAATGC |
|  | Overexpression with P*otef* | sir2StartNcoI | TGCCATGGGCGGCAAAGCCTTCCAAAG |
|  |  | sir2StopNotI | CAGCGGCCGCTTATGATGGAGCCTGTTGTG |
|  | Overexpression with P*otef* | sir2StartSacII | TATCCGCGGATGGGCGGCAAAGCCTTCC |
|  |  | sir2StopXbaI | ATTCTAGATTATGATGGAGCCTGTTGTG |
|  | *ip* locus integration verification | N_Sdh2_fwd | TCCTGTCTTTTCGGCAAGACTCTTCG |
|  |  | N_pDL51_otef_rev | TGGTGCACTCTCAGTACAATCTGC |
|  |  | Amp1_fwd | TTCTGTGACTGGTGAGTACTCAACC |
|  |  | N_Sdh2_rev | TAAGTGACGATTGCGAGTTCTCTTGG |
| *hst2* | Deletion construct | hst2KO5_fwd | GCCATTGTTGTGTGTGTATGGATCG |
|  |  | hst2KO5_rev | CACGGCCTGAGTGGCCGGCATCCAATCCCAGAATACG |
|  |  | hst2KO3_fwd | GTGGCCATCTAGGCCTTTGGTCGTGGTGGTTGTGC |
|  |  | hst2KO3_rev | CGCATACGAGACAGAGACAGG |
|  | Deletion verification | hst2CO5_fwd | TACACCGGCATTGTTAATCAGC |
|  |  | nat2_rev | AGATGATGATGCCGATGCTACC |
|  |  | nat1_fwd | TGGCTGCTGATCACAGCAAGTCAGATT |
|  |  | hst2CO3_rev | GAAAGCTCATTTCTTCCCGTCG |
|  |  | hst2KOint_fwd | TGTATCCGGGCAACTTCAAGC |
|  |  | hst2KOint_rev | CCCATGACAATGAGTAGGTCG |
|  | eGFP tagging | hst2ORF_fwd | CGAATTCCTGCAGCCCGGGGATGCCAGAGACAAAGAGC |
|  |  | hst2ORF_rev | TGCTCACCATTGATGACGGTTTGTTCGAAAC |
|  |  | eGFPHyg_hst2_fwd | ACCGTCATCAATGGTGAGCAAGGGCGAG |
|  |  | eGFPHyg_ hst2_rev | CGGCGTCTCCTATTAATGCGGCCGCACAG |
|  |  | hst2Ter_fwd | CGCATTAATAGGAGACGCCGACATGCAG |
|  |  | hst2Ter_rev | CGGCCGCTCTAGAACTAGTGTGATCTGCTTTCCTCCTTATCCG |
|  |  | hst2GFP_fwd | ATGCCAGAGACAAAGAGC |
|  |  | hst2GFP_rev | TGATCTGCTTTCCTCCTTATCCG |
| *hst4* | Deletion construct | hst4KO5_fwd | AGGATGACAGACAATCCACC |
|  |  | hst4KO5_rev | CACGGCCTGAGTGGCCATGACTCGTGACGAAGTAGG |
|  |  | hst4KO3_fwd | GTGGCCATCTAGGCCTATGCATGATGTGTGTTCG |
|  |  | hst4KO3_rev | TGGGTAGGAAGGTAAGCAGG |
|  | Deletion verification | hst4CO5_fwd | TTCACAGAAAGAGGAAAAGC |
|  |  | gen1_rev | TCTTCTGAGCGGGACTCTGG |
|  |  | gen2_fwd | GTACGGGTACATCGGATCTGC |
|  |  | hst4CO3_rev | CTGGTTTCCAATCTGTCTGG |
|  | eGFP tagging | hst4ORF_fwd | CGAATTCCTGCAGCCCGGGGCCTGCTGAGGCCAAAGCC |
|  |  | hst4ORF_rev | TGCTCACCATAAGACAAGCAGCCGTCTC |
|  |  | eGFPHyg_hst4_fwd | TGCTTGTCTTATGGTGAGCAAGGGCGAG |
|  |  | eGFPHyg_ hst4_rev | CTATATGCGCTATTAATGCGGCCGCACAG |
|  |  | hst4Ter_fwd | CGCATTAATAGCGCATATAGTGCGGAATG |
|  |  | hst4Ter_rev | CGGCCGCTCTAGAACTAGTGGTGTCCTAAGTCGAAATTG |
|  |  | hst4GFP_fwd | CCTGCTGAGGCCAAAGCC |
|  |  | hst4GFP_rev | GTGTCCTAAGTCGAAATTGG |
| *hst5* | Deletion construct | hst5KO5_fwd | GTACTACTGCATGGTCAAGC |
|  |  | hst5KO5_rev | CACGGCCTGAGTGGCCCTAGCCAAGTGCACTCTTGC |
|  |  | hst5KO3_fwd | GTGGCCATCTAGGCCATAGAGGATGAAACCACGCG |
|  |  | hst5KO3_rev | ACCTTGTTGGTCTTTCTTGG |
|  | Deletion verification | hst5CO5_fwd | GCGAAGACGATGAAAATGG |
|  |  | gen1_rev | TCTTCTGAGCGGGACTCTGG |
|  |  | gen2_fwd | GTACGGGTACATCGGATCTGC |
|  |  | hst5CO3_rev | GAAGCTCAGCGAATCATGC |
|  |  | hst5KOint_fwd | AGAAGGCTACCGTAAAGACG |
|  |  | hst5KOint_rev | ACGATTCAGGACCATGACG |
|  | eGFP Tagging | hst5ORF_fwd | GATATCGAATTCCTGCAGCCCGGGGACATTATAGGCCGATCTTCTACCAC |
|  |  | hst5ORF_rev | CCTTGCTCACCATACTGCTGATCACGCCGCT |
|  |  | eGFPHyg_hst5_fwd | CGTGATCAGCAGTATGGTGAGCAAGGGCGAG |
|  |  | eGFPHyg_ hst5_rev | ATCCTCTATGACCTATTAATGCGGCCGCACAG |
|  |  | hst5Ter_fwd | GCCGCATTAATAGGTCATAGAGGATGAAACCACG |
|  |  | hst5Ter_rev | GGTGGCGGCCGCTCTAGAACTAGTGAGTCGCTCCACGCACTTC |
|  |  | hst5GFP_fwd | ACATTATAGGCCGATCTTCTACC |
|  |  | hst5GFP_rev | AGTCGCTCCACGCACTTC |
| *hst6* | Deletion construct | hst6KO5_fwd | GATATCGAATTCCTGCAGCCCGGGGCTACCGTGGGGACAGACAG |
|  |  | hst6KO5_rev | ATGGTGGCCATCTTTCGTTACGAGCATAAGGCAAATG |
|  |  | cbx_hst6KO_fwd | TGCTCGTAACGAAAGATGGCCACCATGGCGT |
|  |  | cbx_hst6KO_rev | GTGATAGTCCGCACTCAGGCCTATTAATGCGGC |
|  |  | hst6KO3_fwd | AATAGGCCTGAGTGCGGACTATCACGTTCTTC |
|  |  | hst6KO3_rev | GGTGGCGGCCGCTCTAGAACTAGTGTAGTCGAGCCATCTGGTG |
|  | Deletion verification | hst6CO5_fwd | CTGCCTGACTCATCATTTGC |
|  |  | cbx1_rev | TCTGGGTTTCGCGAGAGATCTCACAGAGCA |
|  |  | cbx2_fwd | AATTGCACAGATCAAGAAGGACATGGCCGT |
|  |  | hst6CO3_rev | ATTCAAGTTCTCCCAGATGC |
|  |  | hst6KOint_fwd | CAAAGTCTGGCAGTTCTACC |
|  |  | hst6KOint_rev | CTAGCTCTGGAATCGATTCG |
|  | eGFP tagging | hst6ORF_fwd | CGAATTCCTGCAGCCCGGGGGATGTCGACTCTTGCGGCAAACC |
|  |  | hst6ORF_rev | TGCTCACCATAAGGCCAAGCACCTCTGG |
|  |  | eGFPHyg_hst6_fwd | GCTTGGCCTTATGGTGAGCAAGGGCGAG |
|  |  | eGFPHyg_ hst6_rev | CCGCTCAAAGTATTAATGCGGCCGCACAG |
|  |  | hst6Ter_fwd | CGCATTAATACTTTGAGCGGACTATCAC |
|  |  | hst6Ter_rev | CGGCCGCTCTAGAACTAGTGGTCACGACGATGGCAATG |
|  |  | hst6GFP_fwd | GATGTCGACTCTTGCGGCAAACC |
|  |  | hst6GFP_rev | GTCACGACGATGGCAATG |
| Others | RT-qPCR | ppi1_qPCR_fwd | ACATCGTCAAGGCTATCG |
|  |  | ppi1_qPCR_rev | AAAGAACACCGGACTTGG |
|  |  | gapdh_qPCR_fwd | CTTCGGCATTGTTGAGGGTTTG |
|  |  | gapdh_qPCR_rev | TCCTTGGCTGAGGGTCCGTC |
|  |  | sir2_qPCR_fwd | CAAAGTCGCACCTGTATCCGA |
|  |  | sir2_qPCR_rev | AGTGGAGGATTGGCGTCTTGT |
|  |  | eff1-9_qPCR_fwd | CAAGCAAAGAATCCGATCGAGT |
|  |  | ppi1_ChIP_fwd | GGAGGCAAGTCGATCTACGG |
|  |  | ppi1_ChIP_rev | CATGGAAAGAAGACCGGGCT |
|  |  | eff1-9_Pr_ChIP_fwd | ACCTCGCAGCTCAAGGGTAA |
|  |  | eff1-9_Pr_qPCR_rev | AATACCTACCCGCCTGTGAG |
|  |  | eff1-9_orf_ChIP_fwd | AGCAAGCGCGGTGTATGCGA |
|  |  | eff1-9_orf_ChIP_rev | TGCATTTCTCTGACGCTGAG |
|  |  | 06128_Pr_ChIP_fwd | GCCAGGCTGCCAAATAAAAC |
|  |  | 06128_Pr_ChIP_rev | TGGCGCGGAAAAACCAATAC |
|  |  | 06128_orf_ChIP_fwd | TCGCCTGCATCATATTCCAC |
|  |  | 06128_orf_ChIP_rev | TTTCGCGGTTGGAAAACTCG |
|  |  | rsp3_Pr_ChIP_fwd | AGCCTTCTTTTCCACACTGC |
|  |  | rsp3_Pr_ChIP_rev | TGGGGAAACGAGGTTATGAG |
|  |  | rsp3_orf_ChIP_fwd | AGCAAAAGCAAGACGAGCAG |
|  |  | rsp3_orf_ChIP_rev | TCCTTCTTTTGACGCTCCTG |
|  |  | mig2-3_Pr_ChIP_fwd | ATTGTGCGCACATTGCTCTG |
|  |  | mig2-3_Pr_ChIP_rev | TGCTTGAGCTGACTGTATGC |
|  |  | mig2-3_orf_ChIP_fwd | GTTTCCCAGCTTGTTCCTAACG |
|  |  | mig2-3_orf_ChIP_rev | AAAAGCACTGTCCGATAGCG |
|  |  | mig2-6_Pr_ChIP_fwd | TATGATTCTCAGGCGCAGTG |
|  |  | mig2-6_Pr_ChIP_rev | CACCCAAATCTTCCCACATC |
|  |  | mig2-6_orf_ChIP_fwd | AGGATTCGACCATTCTCCAC |
|  |  | mig2-6_orf_ChIP_rev | CAATGTGAACGACAGGCATC |
|  |  | 01241_Pr_ChIP_fwd | TTCTGCCGCATCTAAGTGTG |
|  |  | 01241_Pr_ChIP_rev | AACGGCACAAACAGTACGTC |
|  |  | 01241_orf_ChIP_fwd | ATGTCATGTGGCAGAATCGG |
|  |  | 01241_orf_ChIP_rev | AAGCCAGCCAGGGATTTTTC |

| **Replicate 1** | | | | | | |
| --- | --- | --- | --- | --- | --- | --- |
| **Strain** | **No Tumors** | **Small Tumors** | **Medium Tumors** | **Heavy Tumors** | **Dead Plants** | **n** |
| **WT** | 31 | 37 | 29 | 10 | 5 | 112 |
| **Δ*sir2*** | 17 | 37 | 22 | 13 | 6 | 95 |
| **Replicate 2** | | | | | | |
| **Strain** | **No Tumors** | **Small Tumors** | **Medium Tumors** | **Heavy Tumors** | **Dead Plants** | **n** |
| **WT** | 8 | 13 | 13 | 2 | 2 | 38 |
| **Δ*sir2*** | 3 | 12 | 11 | 6 | 4 | 36 |
| **Replicate 3** | | | | | | |
| **Strain** | **No Tumors** | **Small Tumors** | **Medium Tumors** | **Heavy Tumors** | **Dead Plants** | **n** |
| **WT** | 3 | 6 | 13 | 4 | 3 | 29 |
| **Δ*sir2*** | 3 | 7 | 11 | 4 | 4 | 29 |
| **Replicate 1 + Replicate 2 + Replicate 3** | | | | | | |
| **Strain** | **No Tumors** | **Small Tumors** | **Medium Tumors** | **Heavy Tumors** | **Dead Plants** | **n** |
| **WT** | 42 | 56 | 55 | 16 | 10 | 179 |
| **Δ*sir2*** | 23 | 56 | 44 | 23 | 14 | 160 |
| **Percentage** | | | | | |  |
| **Strain** | **No Tumors** | **Small Tumors** | **Medium Tumors** | **Heavy Tumors** | **Dead Plants** |  |
| **WT** | 23,46 | 31,28 | 30,73 | 8,94 | 5,59 |  |
| **Δ*sir2*** | 14,38 | 35,00 | 27,50 | 14,38 | 8,75 |  |

**TABLE S3. Individual infection data of the indicated strains.**

| **Replicate 1** | | | | | | |
| --- | --- | --- | --- | --- | --- | --- |
| **Strain** | **No Tumors** | **Small Tumors** | **Medium Tumors** | **Heavy Tumors** | **Dead Plants** | **n** |
| **WT** | 9 | 3 | 4 | 4 | 1 | 21 |
| **Δ*hst2*** | 8 | 5 | 1 | 8 | 1 | 23 |
| **Replicate 2** | | | | | | |
| **Strain** | **No Tumors** | **Small Tumors** | **Medium Tumors** | **Heavy Tumors** | **Dead Plants** | **n** |
| **WT** | 6 | 7 | 7 | 21 | 5 | 46 |
| **Δ*hst2*** | 12 | 4 | 6 | 20 | 7 | 49 |
| **Replicate 1 + Replicate 2** | | | | | | |
| **Strain** | **No Tumors** | **Small Tumors** | **Medium Tumors** | **Heavy Tumors** | **Dead Plants** | **n** |
| **WT** | 15 | 10 | 11 | 25 | 6 | 67 |
| **Δ*hst2*** | 20 | 9 | 7 | 28 | 8 | 72 |

| **Percentage** | | | | | |
| --- | --- | --- | --- | --- | --- |
| **Strain** | **No Tumors** | **Small Tumors** | **Medium Tumors** | **Heavy Tumors** | **Dead Plants** |
| **WT** | 22,39 | 14,93 | 16,42 | 37,31 | 8,96 |
| **Δ*hst2*** | 27,78 | 12,50 | 9,72 | 38,89 | 11,11 |

| **Replicate 1** | | | | | | |
| --- | --- | --- | --- | --- | --- | --- |
| **Strain** | **No Tumors** | **Small Tumors** | **Medium Tumors** | **Heavy Tumors** | **Dead Plants** | **n** |
| **WT** | 10 | 3 | 4 | 8 | 2 | 27 |
| **Δ*hst5*** | 9 | 1 | 3 | 8 | 2 | 23 |
| **Δ*hst6*** | 3 | 4 | 4 | 11 | 2 | 24 |
| **Replicate 2** | | | | | | |
| **Strain** | **No Tumors** | **Small Tumors** | **Medium Tumors** | **Heavy Tumors** | **Dead Plants** | **n** |
| **WT** | 6 | 7 | 7 | 21 | 5 | 46 |
| **Δ*hst5*** | 7 | 9 | 6 | 15 | 6 | 43 |
| **Δ*hst6*** | 7 | 10 | 3 | 13 | 8 | 41 |
| **Replicate 1 + Replicate 2** | | | | | | |
| **Strain** | **No Tumors** | **Small Tumors** | **Medium Tumors** | **Heavy Tumors** | **Dead Plants** | **n** |
| **WT** | 16 | 10 | 11 | 29 | 7 | 73 |
| **Δ*hst5*** | 16 | 10 | 9 | 23 | 8 | 66 |
| **Δ*hst6*** | 10 | 14 | 7 | 24 | 10 | 65 |
| **Percentage** | | | | | |  |
| **Strain** | **No Tumors** | **Small Tumors** | **Medium Tumors** | **Heavy Tumors** | **Dead Plants** |  |
| **WT** | 21,92 | 13,70 | 15,07 | 39,73 | 9,59 |  |
| **Δ*hst5*** | 24,24 | 15,15 | 13,64 | 34,85 | 12,12 |  |
| **Δ*hst6*** | 15,38 | 21,54 | 10,77 | 36,92 | 15,38 |  |

| **Replicate 1** | | | | | | |
| --- | --- | --- | --- | --- | --- | --- |
| **Strain** | **No tumors** | **Small Tumors** | **Medium Tumors** | **Heavy Tumors** | **Dead Plants** | **n** |
| **WT** | 4 | 4 | 2 | 5 | 4 | 19 |
| **P*pit2*:*sir2* 1c** | 7 | 5 | 2 | 4 | 5 | 23 |
| **Replicate 2** | | | | | | |
| **Strain** | **No tumors** | **Small Tumors** | **Medium Tumors** | **Heavy Tumors** | **Dead Plants** | **n** |
| **WT** | 9 | 5 | 2 | 4 | 2 | 22 |
| **P*pit2*:*sir2* 1c** | 5 | 5 | 3 | 7 | 3 | 23 |
| **Replicate 1 + Replicate 2** | | | | | | |
| **Strain** | **No Tumors** | **Small Tumors** | **Medium Tumors** | **Heavy Tumors** | **Dead Plants** | **n** |
| **WT** | 13 | 9 | 4 | 9 | 6 | 41 |
| **P*pit2*:*sir2* 1c** | 12 | 10 | 5 | 11 | 8 | 46 |
| **Percentage** | | | | | |  |
| **Strain** | **No Tumors** | **Small Tumors** | **Medium Tumors** | **Heavy Tumors** | **Dead Plants** |  |
| **WT** | 30,00 | 21,43 | 11,43 | 24,29 | 12,86 |  |
| **P*pit2*:*sir2* 1c** | 26,09 | 21,74 | 10,87 | 23,91 | 17,39 |  |

| **Replicate 1** | | | | | | |
| --- | --- | --- | --- | --- | --- | --- |
| **Strain** | **No tumors** | **Small Tumors** | **Medium Tumors** | **Heavy Tumors** | **Dead Plants** | **n** |
| **WT** | 14 | 11 | 8 | 19 | 1 | 53 |
| **P*pit2*:*sir2* >1c** | 47 | 3 | 0 | 0 | 0 | 50 |
| **Replicate 2** | | | | | | |
| **Strain** | **No tumors** | **Small Tumors** | **Medium Tumors** | **Heavy Tumors** | **Dead Plants** | **n** |
| **WT** | 13 | 3 | 10 | 13 | 1 | 40 |
| **P*pit2*:*sir2* >1c** | 36 | 14 | 0 | 0 | 0 | 50 |
| **Replicate 3** | | | | | | |
| **Strain** | **No tumors** | **Small Tumors** | **Medium Tumors** | **Heavy Tumors** | **Dead Plants** | **n** |
| **WT** | 12 | 7 | 4 | 9 | 0 | 32 |
| **P*pit2*:*sir2* >1c** | 22 | 12 | 0 | 0 | 0 | 34 |
| **Replicate 1 + Replicate 2 + Replicate 3** | | | | | | |
| **Strain** | **No Tumors** | **Small Tumors** | **Medium Tumors** | **Heavy Tumors** | **Dead Plants** | **n** |
| **WT** | 39 | 21 | 22 | 41 | 2 | 125 |
| **P*pit2*:*sir2* >1c** | 105 | 29 | 0 | 0 | 0 | 134 |
| **Percentage** | | | | | |  |
| **Strain** | **No Tumors** | **Small Tumors** | **Medium Tumors** | **Heavy Tumors** | **Dead Plants** |  |
| **WT** | 31,20 | 16,80 | 17,60 | 32,80 | 1,60 |  |
| **P*pit2*:*sir2* >1c** | 78,36 | 21,64 | 0,00 | 0,00 | 0,00 |  |
